# RNA decay via the nuclear exosome is essential for Piwi-mediated transposon silencing

**DOI:** 10.64898/2025.12.16.694471

**Authors:** Changwei Yu, Toni Manolova, Laszlo Tirian, Dominik Handler, Ulrich Hohmann, Filip Nemčko, Júlia Portell-Montserrat, Guillaume Giraud Collet, Jakob Schnabl-Baumgartner, Ellie Blake, Peter Duchek, Maria Novatchkova, Elisabeth Roitinger, Felipe Karam Teixeira, Sebastian Falk, Julius Brennecke

**Affiliations:** Institute of Molecular Biotechnology of the Austrian Academy of Sciences (IMBA), Vienna BioCenter (VBC); Vienna, 1030, Austria; Max Perutz Labs, Vienna Biocenter Campus (VBC); Vienna, 1030, Austria; University of Vienna, Max Perutz Labs, Department of Structural and Computational Biology; Vienna, 1030, Austria; Vienna BioCenter PhD Program, Doctoral School of the University of Vienna and Medical University of Vienna; Vienna, 1030, Austria; Research Institute of Molecular Pathology (IMP), Vienna BioCenter (VBC); Vienna, 1030, Austria; Université Paris Cité, Magistère Européen de génétique; Paris, 75006, France; University of Cambridge, Department of Genetics, Cambridge CB2 3EH, UK; University of Cambridge, Department of Physiology, Development and Neuroscience, Cambridge CB2 3DY, UK

## Abstract

Nuclear Argonaute proteins safeguard genome integrity by directing transcriptional silencing and heterochromatin formation at transposon loci. Yet it remains unclear how Argonautes enforce robust repression while relying on target transcription for their own recruitment. Here we show that transposon silencing by the *Drosophila* nuclear Piwi–piRNA pathway requires degradation of target RNA by the nuclear exosome. Using proximity proteomics at endogenous Piwi target sites, we identify two previously uncharacterized paralogs, TEsup-1 and TEsup-2, as essential cofactors for Piwi-mediated silencing. TEsup proteins act in part by engaging nuclear exosome adaptor complexes at piRNA-targeted transcripts through a domain that recognizes proline-rich peptides. Disruption of the Piwi–TEsup–exosome axis leads to accumulation and nuclear export of piRNA-targeted transposon RNAs. Notably, the *P*-element—which evades heterochromatin-based repression—is silenced primarily through this RNA-decay pathway. Thus, the nuclear piRNA pathway couples target recognition to RNA degradation, reconciling small RNA–guided heterochromatin formation with ongoing transcription at target loci.

## INTRODUCTION

Genome integrity is under constant threat from transposable elements (TEs), ubiquitous selfish genetic elements with high mutagenic potential (Bourque *et al*, 2018; Fedoroff, 2012). Locked in an evolutionary arms race, organisms have evolved diverse mechanisms to restrict TE activity (Levin & Moran, 2011). A deeply conserved defense strategy employs nuclear small RNA pathways, which use small regulatory RNAs to guide Argonaute proteins to complementary nascent transcripts, triggering transcriptional repression and heterochromatin formation (Grewal, 2010; Holoch & Moazed, 2015; Martienssen & Moazed, 2015). This mechanism, however, creates a fundamental paradox: Argonaute recruitment requires active transcription of the very target loci being silenced. How cells achieve efficient repression while relying on ongoing target transcription remains poorly understood.

Pioneering studies in fungi, plants, and animals have greatly advanced our understanding of how nuclear small RNA pathways orchestrate heterochromatin-based repression (Czech & Hannon, 2016; Grewal, 2010; Holoch & Moazed, 2015; Martienssen & Moazed, 2015; Ozata *et al*, 2018). By contrast, it remains unclear whether these pathways also control the fate of their target RNAs—for example by promoting degradation, inducing transcription termination, or restricting nuclear export—and whether such RNA-level processes operate alongside heterochromatin formation to help reconcile the paradox of transcription-dependent silencing.

The *Drosophila* PIWI-interacting RNA (piRNA) pathway in ovarian follicle cells provides a powerful system to dissect how the nuclear Argonaute protein Piwi acts on its nascent target RNAs. In this pathway, the genomic loci that generate piRNAs are spatially distinct from the TE insertions they silence, enabling the analysis of transcriptional repression independently of small RNA biogenesis (Czech *et al*, 2018; Ozata *et al*., 2018; Senti & Brennecke, 2010; Sienski *et al*, 2012). Piwi-dependent silencing relies on several pathway-specific cofactors as well as the H3K9-methylation, H3K4-demethylation, and SUMOylation machineries (Andreev *et al*, 2022; Batki *et al*, 2019; Donertas *et al*, 2013; Eastwood *et al*, 2021; Fabry *et al*, 2019; Le Thomas *et al*, 2013; Muerdter *et al*, 2013; Mugat *et al*, 2020; Murano *et al*, 2019; Ninova *et al*, 2020; Ohtani *et al*, 2013; Rozhkov *et al*, 2013; Schnabl *et al*, 2021; Sienski *et al*., 2012; Wang & Elgin, 2011; Yang *et al*, 2019). Upon binding a piRNA-complementary transcript, Piwi forms an activated complex, Piwi*, composed of Piwi, Maelstrom, and Asterix/Gtsf1 (De *et al*, 2025; Portell-Montserrat *et al*, 2025). Through composite protein interfaces, Piwi* engages downstream effectors such as the SFiNX repressor complex to active TE insertions (Portell-Montserrat *et al*., 2025). However, piRNA-guided transcriptional repression precedes heterochromatin establishment, in part through unknown factors (Wu *et al*, 2025), and Piwi has been implicated in the suppression of *P*-element splicing (Teixeira *et al*, 2017), indicating that essential components of the silencing machinery remain to be identified.

Here we identify two previously uncharacterized proteins as critical Piwi cofactors. While both proteins are critical for transcriptional silencing, our work reveals that they further engage the nuclear RNA exosome at piRNA-targeted transcripts, thereby coupling Piwi to RNA decay. This pathway adds an unrecognized layer to nuclear piRNA-mediated repression and resolves how Piwi achieves effective silencing despite ongoing transcription at its target loci.

## RESULTS

### Proximity proteomics at piRNA target sites identifies new Piwi silencing cofactors

To identify proteins that act with Piwi at piRNA target sites, we performed TurboID-based proximity labeling in ovarian somatic cells (OSCs), a well-characterized system with an active nuclear piRNA pathway centered on Piwi (Saito *et al*, 2009; Sienski *et al*., 2012). We generated clonal OSC lines expressing endogenous TurboID fusions (Branon *et al*, 2018) to Piwi, the Piwi* subunit Maelstrom, and the SFiNX subunit Panoramix (Fig. 1A; Fig. S1A). A nuclear TurboID-GFP line served as control (Fig. S1A). Biotinylated proteins were enriched using streptavidin affinity purification and identified by quantitative mass spectrometry (Fig. 1B; Fig. S1B-D).

**Figure 1.**
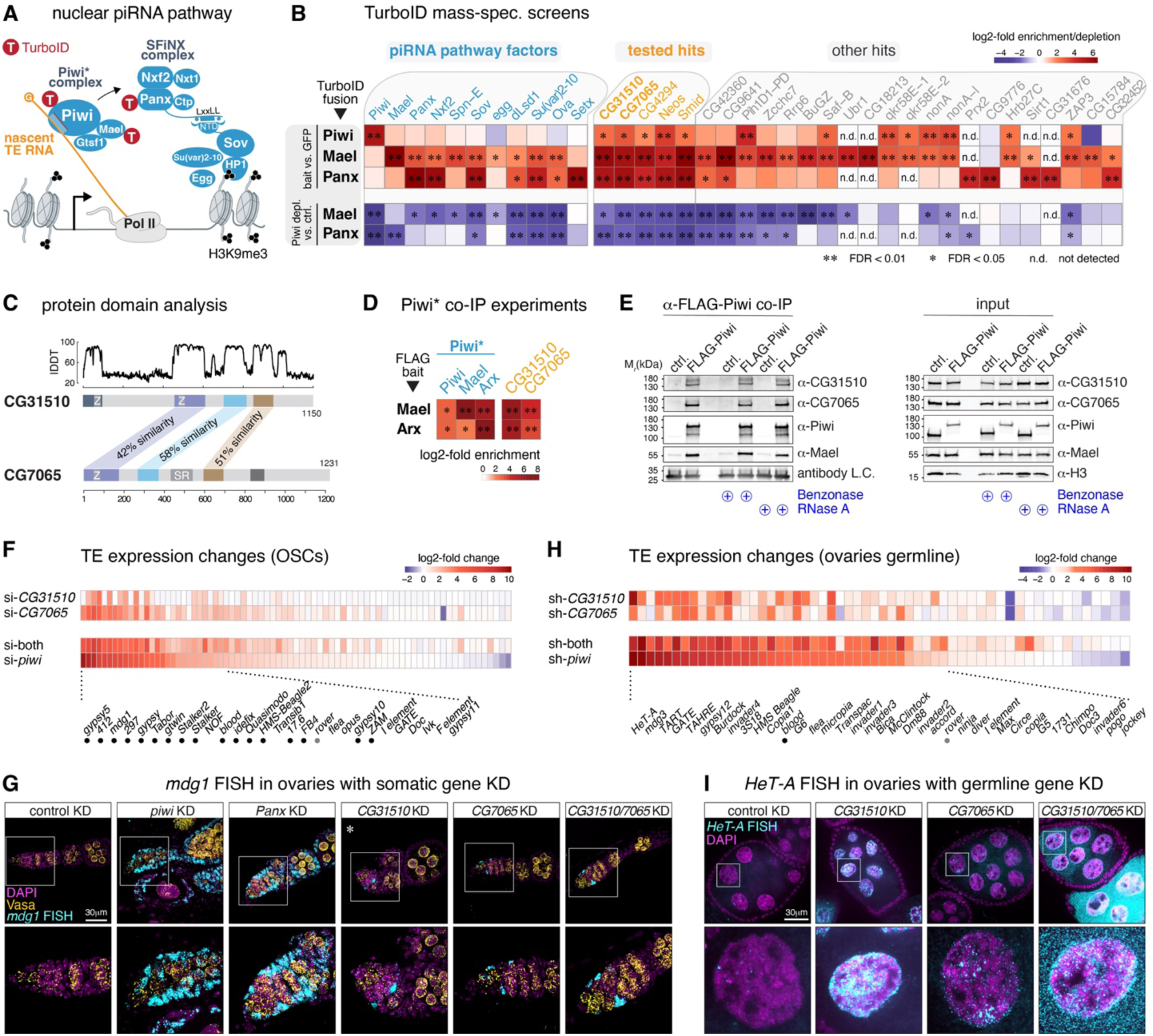
CG31510 and CG7065 are cofactors of Piwi-mediated TE silencing. **(A)** Schematic of the nuclear Piwi–piRNA pathway highlighting TurboID-tagged factors (T) used in proximity biotinylation screens. **(B)** Heatmaps showing protein enrichments in Piwi-, Maelstrom-, and Panoramix-TurboID screens (Table S1) relative to GFP-TurboID control (top), and protein depletion in Maelstrom-and Panoramix-TurboID screens in wild-type versus Piwi-depleted cells (bottom). Known piRNA pathway factors are shown in blue, newly tested factors in orange, and others in grey. Values represent means of three biological replicates (*FDR < 0.05, **FDR < 0.01 based on limma (Hein *et al*, 2015; Smyth, 2004)). **(C)** Domain organization of CG31510 and CG7065 showing structured and unstructured regions (AlphaFold2-predicted local Distance Difference Test, lDDT) and homologous domains with percent similarity indicated (Z, U1-like zinc finger; SR, serine/arginine-rich region). Detailed sequence analysis in Materials & Methods. **(D)** Heatmap showing enrichment of indicated proteins in Maelstrom and Asterix FLAG co-IPs. Values represent means of three biological replicates (*FDR < 0.05, **FDR < 0.01 based on limma). **(E)** Western blots showing specific co-IP of CG31510, CG7065, and Maelstrom with FLAG-Piwi using standard or nuclease-treated OSC lysates as indicated (L.C., antibody light chain). Right panels show input lysates. **(F)** Heatmap of log₂ fold changes in TE RNA levels (poly(A⁺) RNA-seq) following siRNA-mediated depletion of CG31510, CG7065, both, or Piwi in OSCs (n = 3 biological replicates). GeTMM ≥ 5 in at least one dataset, siPiwi RNA-seq data is from (Andreev *et al*., 2022). Dots indicate TEs with known infectivity traits. **(G)** Confocal images of early oogenesis (germaria and stage 2 egg chambers) stained for DNA (DAPI), Vasa, and *mdg1* RNA (FISH) (* indicates weaker FISH signal compared to other conditions). Enlarged views of boxed regions are shown below. **(H)** Heatmap of log₂ fold changes in TE RNA levels (poly(A⁺) RNA-seq) after shRNA-mediated depletion of CG31510, CG7065, both, or Piwi in the ovarian germline (n = 3 biological replicates). GeTMM ≥ 5 in at least one dataset. Dots indicate TEs with known infectivity traits. **(I)** Confocal images of egg chambers stained for DNA (DAPI) and *HeT-A* RNA (FISH). Enlarged views of boxed regions are shown below.

Because only a small fraction of nuclear Piwi is target-engaged at any time (Portell-Montserrat *et al*., 2025), we reasoned that Maelstrom and Panoramix, both of which bind specifically to target-engaged Piwi in the nucleus, would be more effective baits for detecting proteins involved in silencing at piRNA targets. To assess the Piwi dependency of their interactions, we also profiled Maelstrom and Panoramix proximity proteomes in Piwi-depleted cells (Fig. 1B; Fig. S1E,F).

We first validated the TurboID approach by examining established factors in Piwi-mediated silencing. Known cofactors and downstream effectors—including components of the SFiNX complex (Panoramix, Nxf2), the silencing factor Sov, the H3K4 demethylase Lsd1, and the SUMO E3 ligase Su(var)2-10—were significantly enriched (FDR < 0.05) in the Maelstrom and/or Panoramix datasets (Fig. 1B; Fig. S1B–D). In Piwi-depleted cells, where piRNA-targeted TEs are strongly derepressed, enrichment of these factors was lost from the Maelstrom and Panoramix datasets (Fig. 1B; Fig. S1E,F), confirming that TurboID captures bona fide Piwi-dependent interactions. As expected, enrichments in the Piwi dataset were modest (Fig. 1B; Fig. S1B), consistent with most nuclear Piwi not being target-engaged (Portell-Montserrat *et al*., 2025).

In addition to known Piwi cofactors, several proteins were significantly enriched (FDR < 0.05) across the datasets, most showing Piwi-dependent association in Maelstrom and Panoramix proximity proteomes (Fig. 1B). siRNA-mediated depletion of the top five candidates in OSCs (CG31510, CG7065, CG4294, Neos, Smid) using two independent siRNAs led to significant derepression of the Piwi-regulated transposons *mdg1* and *gypsy*, albeit to varying extents (Fig. S1G,H).

Two candidates, CG31510 and CG7065, stood out because they are predicted paralogs (Hu *et al*, 2011), share substantial sequence and domain-architecture similarity (Fig. 1C), and show stronger derepression upon co-depletion than upon individual knockdown (Fig. S2A,B). To test whether both proteins associate with the Piwi* complex, we performed co-IP mass spectrometry with FLAG-Maelstrom and FLAG-Asterix/Gtsf1 (Portell-Montserrat *et al*., 2025). Both proteins were significantly enriched in these co-IPs (Fig. 1D; n = 3; FDR < 0.01). Moreover, CG31510 and CG7065 co-immunoprecipitated with FLAG-Piwi to a similar extent as Maelstrom, and these interactions persisted after RNase A or Benzonase treatment, indicating that their physical association with Piwi is not mediated by nucleic acids (Fig. 1E). Together, these data map the proteome in proximity to Piwi at piRNA target sites and identify CG31510 and CG7065 as previously uncharacterized candidate cofactors in Piwi-mediated silencing.

### TEsup-1 and TEsup-2 are essential cofactors for Piwi-mediated transposon silencing

To define the cellular roles of CG31510 and CG7065, we first analyzed their function in OSCs, where Piwi directly represses a broad set of long terminal repeat (LTR) retrotransposons. siRNA-mediated knockdown of either gene efficiently depleted the target protein and partially reduced its paralog, despite unchanged mRNA levels (Fig. S2A,C). Transcriptome profiling revealed that loss of either factor caused de-repression of multiple Piwi-targeted TE families, and their combined depletion yielded a TE expression pattern that was nearly indistinguishable from that of Piwi knockdown (Pearson *r* = 0.91; Fig. 1F, Fig. S2D). Importantly, host gene expression remained unaffected except for ∼100 genes located adjacent to piRNA-targeted TE insertions, a signature of Piwi-dependent heterochromatin spreading (Fig. S2E) (Sienski *et al*., 2012).

We next examined the function of CG31510 and CG7065 *in vivo*. Endogenous GFP fusions of both proteins localized to the nucleus in somatic and germline cells of developing ovaries (Fig. S3A,B). Tissue-specific RNAi revealed an essential role for both factors in TE repression. In somatic cells, the *mdg1* and *quasimodo* retrotransposons were fully silenced in wild-type ovaries and broadly de-repressed upon Piwi or Panoramix depletion (Fig. 1G; Fig. S3C). Depletion of CG31510 or CG7065 also triggered robust *mdg1* and *quasimodo* expression, but confined to discrete cells in the germarium, corresponding to escort or somatic stem cells, the immediate precursors of OSCs (Fig. 1G; Fig. S3C). This restricted phenotype matches the strict requirement for both factors in OSCs. Quantitatively, >60% of germaria depleted for either paralog and nearly all with double depletion contained strongly TE-positive cells (Fig. S3D).

In the germline, depletion of either factor induced de-repression of numerous transposon families, and combined depletion produced a transposon expression profile that again closely mirrored Piwi loss (Pearson *r* = 0.85; Fig. 1H; Fig. S3E,F). Notably, although the sets of TEs targeted by Piwi in somatic and germline tissues are largely distinct (Fig. 1F,H), CG31510 and CG7065 were required in both contexts to repress the respective Piwi-regulated elements.

Because CG7065 depletion and the double knockdown disrupted ovarian morphology (Fig. 3G), we confirmed germline TE de-repression by RNA FISH for the LINE *HeT-A* and the LTR element *blood* in stage-matched egg chambers (Fig. 1I; Fig. 3H). Consistent with impaired germline transposon silencing, knockdown of either factor reduced female fertility (CG31510: partial sterility; CG7065: complete sterility) (Fig. S3I).

To exclude that the observed phenotypes upon loss of CG31510/CG7065 are due to defective piRNA biogenesis, we analyzed small RNAs and PIWI localization. In OSCs, single or combined depletion of both factors left piRNA levels and size distributions unchanged (Fig. S4A,B). In ovaries, germline depletion of either factor did not alter Piwi’s nuclear localization (Fig. S4C) or overall piRNA abundance and length profiles except for a handful of TEs (Fig. S4D,E). Thus, neither protein affects the production, loading, or nuclear import of Piwi-piRNA complexes.

Together, these results show that CG31510 and CG7065 are indispensable for the silencing of Piwi-targeted transposons in OSCs and flies, establishing them as novel cofactors in Piwi-mediated silencing. Based on their cooperative yet non-redundant roles in repressing transposons, we named them ‘TEs are up-1’ (TEsup-1; CG31510) and ‘TEs are up-2’ (TEsup-2; CG7065).

### TEsup proteins are required for Piwi-directed heterochromatin formation and transcriptional repression

Predicted structures of TEsup-1 and TEsup-2 show no similarity to known proteins and lack recognizable functional domains. To gain insight into their molecular roles, we mapped their protein interactomes using OSC lines expressing endogenous TurboID fusions (Fig. 5A-C). Both proteins exhibited highly overlapping interaction profiles (Fig. 2A), which we validated by co-IP mass spectrometry experiments using FLAG-tagged lines (Fig. 2B; Fig. S5D-F). Three principal features emerged. First, TEsup-1 and TEsup-2 strongly enriched each other, consistent with physical association (Fig. 2B). Second, both proteins co-enriched with core components of the nuclear Piwi silencing machinery, including Piwi, Maelstrom, Asterix/Gtsf1, the SFiNX complex, and the heterochromatin factor Sov (Fig. 2B). Third, in addition to these established silencing factors, their interactomes were prominently enriched for proteins implicated in mRNA metabolic processes (Fig. 2C).

**Figure 2.**
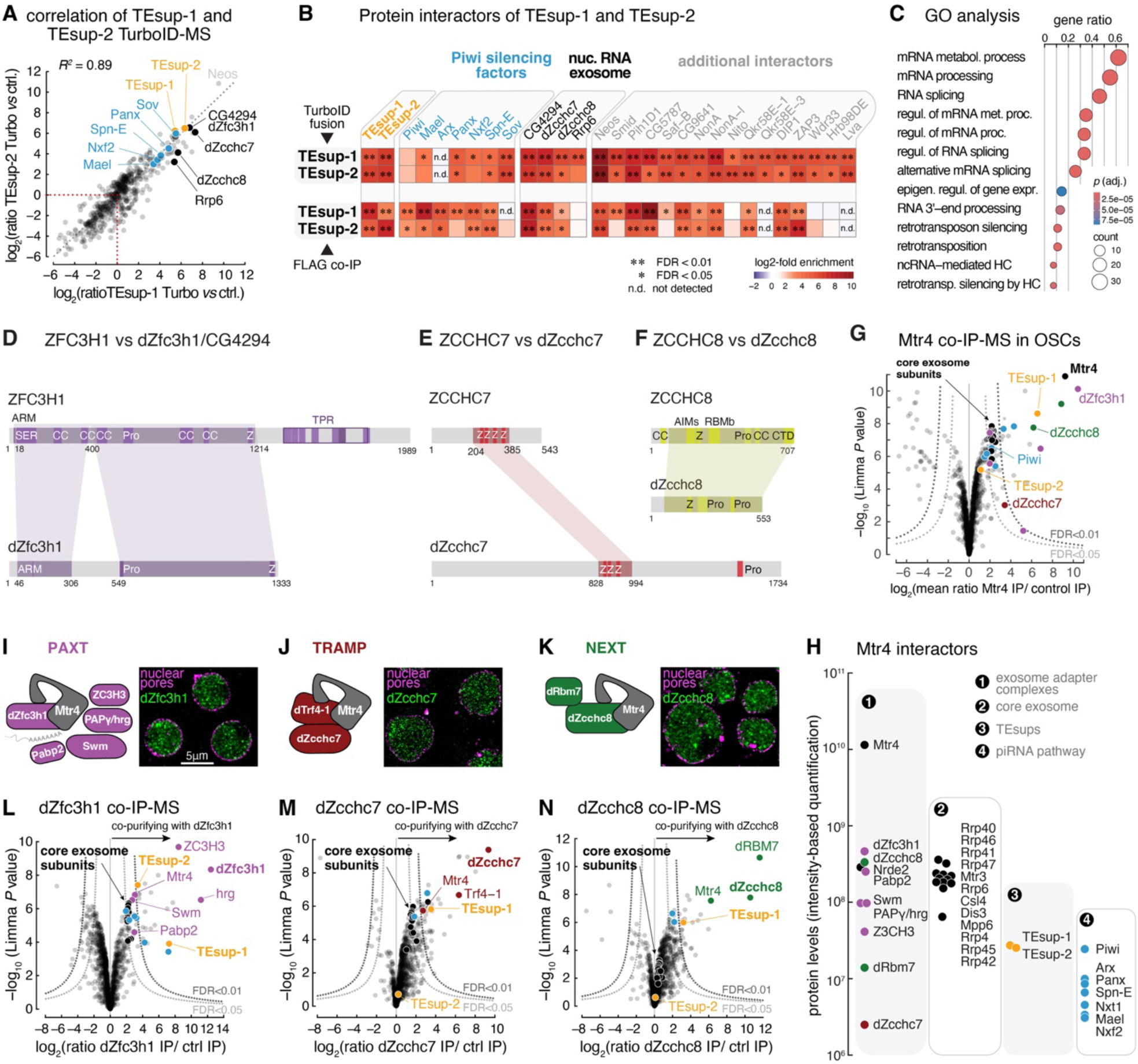
TEsup-1 and TEsup-2 interact with nuclear RNA exosome adaptor complexes. **(A)** Correlation of TEsup-1 and TEsup-2 TurboID experiments (log₂-fold enrichment of proteins versus GFP-TurboID control). Piwi cofactors are shown in blue; exosome-related factors in black. **(B)** Heatmap showing log₂-fold enrichments and FDR categories (**FDR < 0.01; *FDR < 0.05; n.d., not detected) of Piwi silencing factors (blue) and top-enriched proteins in TEsup-1 and TEsup-2 TurboID screens (top; Table S1), with corresponding enrichments in TEsup-1 and TEsup-2 co-IPs (bottom; Table S1). **(C)** Gene Ontology (GO; biological process) enrichment analysis of factors significantly enriched in TEsup-1 or TEsup-2 TurboID datasets (FDR < 0.05; n = 63). Significantly enriched biological processes and associated protein counts are indicated. P values adjusted for multiple testing using the Benjamini–Hochberg method. Abbreviation: HC, heterochromatin formation. **(D)** Schematic representation of human ZFC3H1 and *Drosophila* dZfc3h1 domain organization, with homologous regions indicated. Domain schematics were adapted from (Polak *et al*, 2023) and UniProt protein domains. Similarity analysis was as described in the Material and methods section “Ortholog identification and sequence analysis of CG4294”. Abbreviations: ARM, ARS2 recruitment motif; SER, serine-rich; CC, coiled coil; PRO, proline-rich; C_3_H_1_ type Zinc finger; TPR, tetratricopeptide repeats. **(E)** Schematic representation of human ZCCHC7 and *Drosophila* dZcchc7 domain organization, with homologous regions indicated. Domain schematics were adapted from UniProt and SMART protein domains. Homology prediction was determined by HHpred (Soding *et al*, 2005). Abbreviations: Z, C_2_H_1_ type Zinc finger; Pro, Pro-rich domain. **(F)** Schematic representation of human ZCCHC8 and *Drosophila* dZcchc8 domain organization, with homologous regions indicated. Domain organization was adapted from (Gerlach *et al*, 2022) and Pfam protein domains. Homology prediction was determined by HHpred. Abbreviations: CC, coiled coil; Z, Zinc finger; AIMs, arch-interacting motifs; RBMb, RBM binding domain; Pro, Pro-rich domain; CTD, C-terminal domain. **(G)** Volcano plot showing log2-fold enrichment and significance of proteins identified by quantitative mass spectrometry in Mtr4 co-IP versus control (n = 3 biological replicates). Selected factors are labeled: nuclear RNA exosome adapter complex PAXT co-factors (purple), TRAMP subunits (red), NEXT subunits (green); RNA exosome core (black); TEsup-1 and TEsup-2 (orange); nuclear piRNA pathway factors (blue). **(H)** Protein abundances of Mtr4 interactors identified by quantitative mass spectrometry in Mtr4 co-IPs versus control (n = 3 biological replicates). Shown are (1) nuclear RNA exosome adaptor complex factors of PAXT (purple), TRAMP (red), and NEXT (green); (2) RNA exosome subunits (black); (3) TEsup proteins (orange); and (4) Piwi silencing factors (blue). **(I–K)** Confocal images showing localization of FLAG–GFP–tagged dZfc3h1 (I), dZcchc7 (J), and dZcchc8 (K) in OSCs (GFP, green; nuclear pores (WGA), magenta; scale bar, 5 µm) alongside cartoons of the three corresponding exosome adaptor complexes. **(L–N)** Volcano plots showing proteins co-purifying with dZfc3h1 (L), dZcchc7 (M), or dZcchc8 (N) in co-IP experiments versus control (n = 3 biological replicates; Table S1). Color code as in (I), Piwi cofactors shown as blue and exosome subunits as black dots.

To determine whether TEsup proteins function directly within the Piwi-directed silencing pathway, we examined their contribution to heterochromatin formation and transcriptional repression at endogenous targets. Depletion of TEsup-2, and to a lesser extent TEsup-1, reduced H3K9me3 levels and increased RNA polymerase II occupancy at the 381 previously defined piRNA-targeted TE insertions in OSCs (Fig. S6A–D). Co-depletion of both paralogs resulted in defects in heterochromatin formation and transcriptional repression that closely matched those observed upon Piwi loss (Fig. S6A–D). Because these analyses rely exclusively on uniquely mapped reads flanking individual TE insertions, they demonstrate that Piwi and TEsup proteins act at the same genomic loci, providing strong evidence for functional cooperation.

Consistent with this conclusion, experimental tethering of either TEsup-1 or TEsup-2 to a nascent reporter RNA using the λN–boxB system induced robust transcriptional repression (Fig. S6E,F). In each case, silencing was accompanied by local H3K9 trimethylation at the reporter locus in OSCs (Fig. S6G).

Together, these findings establish that TEsup-1 and TEsup-2 function downstream of Piwi in piRNA-guided transposon silencing. Notably, however, their strong GO-term association with factors involved in mRNA metabolism (Fig. 2C) suggests that their role in the piRNA pathway may extend beyond chromatin-based repression, pointing to an additional layer of regulation that acts at the level of RNA.

### TEsup proteins interact with nuclear RNA exosome adaptor complexes

To explore whether the association of TEsup proteins with RNA metabolic factors reflects a specific functional connection, we examined their interactomes in greater detail. Across both TurboID proximity labeling and co-immunoprecipitation datasets for TEsup-1 and TEsup-2, three proteins—CG4294, dZcchc7, and CG4622—consistently ranked among the most strongly enriched interactors (Fig. 2B). Phylogenetic analysis identified these factors as the *Drosophila* homologs of human ZFC3H1, ZCCHC7, and ZCCHC8, which serve as scaffold subunits of the PAXT, TRAMP, and NEXT adaptor complexes that deliver RNA substrates to the nuclear exosome for degradation or 3′ processing (Fig. 2D–F) (Januszyk & Lima, 2014; Schmid & Jensen, 2019).

The nuclear exosome is a conserved 3′–5′ exoribonuclease complex that relies on adaptor systems, each centered on a distinct scaffold protein and the shared RNA helicase Mtr4, to channel specific RNA substrates to its nuclease subunits (Januszyk & Lima, 2014; Schmid & Jensen, 2019). Although these adaptor complexes have been extensively characterized in yeast and mammals, their composition and function in *Drosophila* remain only partially understood (Eberle *et al*, 2015; Lee *et al*, 2025). We therefore sought to systematically characterize nuclear exosome adaptor complexes in OSCs.

Endogenous co-IP mass-spectrometry experiments using FLAG-tagged Mtr4, common to all adaptor systems, recovered the three scaffold proteins dZfc3h1 (CG4294), dZcchc7, and dZcchc8 (CG4622), together with signature components of their respective adaptor complexes (Fig. 2G,H; Fig. S7A). Endogenous FLAG-GFP fusions of each scaffold localized to the nucleus (Fig. 2I–K; Fig. S7A) and robustly interacted with Mtr4 (Fig. 2L–N). Analysis of their individual interactomes further confirmed adaptor identity: dZfc3h1 associated with known PAXT components (Pabp2, Swm, dZC3H3, Hrg), dZcchc7 with the TRAMP-specific poly(A) polymerase Trf4, and dZcchc8 with the NEXT-specific RNA-binding protein dRbm7 (CG11454) (Fig. 2L–N).

Functionally, the *Drosophila* adaptors recapitulated key features of their mammalian counterparts. siRNA-mediated depletion of individual adaptors followed by transient transcriptome sequencing (TT-seq) in OSCs revealed distinct RNA-processing defects. Loss of dZcchc7 (TRAMP) impaired 3′ processing of snRNAs and snoRNAs (Fig. S7B,C), whereas depletion of dZcchc8 (NEXT) led to accumulation of Drosha-processed microRNA precursors as well as snRNAs and snoRNAs (Fig. S7B–D). In contrast, depletion of dZfc3h1 (PAXT), but not NEXT, resulted in elevated levels of cellular transcripts that increase in core exosome-depleted cells (Fig. S7E) (Hrossova *et al*, 2015; Imamura *et al*, 2024; Lee *et al*., 2025; Lubas *et al*, 2015). Thus, *Drosophila* harbors functional analogs of the human PAXT, TRAMP and NEXT adaptor complexes.

Notably, TEsup proteins, particularly TEsup-1, along with key Piwi silencing factors including Piwi, Maelstrom, Asterix, the SFiNX complex, were significantly enriched in co-IPs of Mtr4 and the adaptor scaffolds, most prominently with dZfc3h1 (Fig. 2G,H, 2L–N). These data uncover a physical association between nuclear exosome adaptor complexes and the Piwi silencing machinery, suggesting that TEsup proteins may serve as molecular links connecting Piwi-directed target recognition to nuclear RNA degradation.

### A proline-recognition domain enables TEsup proteins to recruit RNA exosome adaptors

To determine whether TEsup proteins engage nuclear exosome adaptor complexes through direct molecular interactions, we used AlphaFold2-multimer to predict interactions between TEsup proteins and candidates from their interactomes (Evans *et al*, 2022; Jumper *et al*, 2021). This analysis identified a conserved α-helical region in both TEsup proteins that formed high-confidence interaction interfaces (PEAK score > 0.75; iPTM ≈ 0.75) with the PAXT scaffold dZfc3h1, as well as with the Piwi-associated factors Pih1D1 and Saf-B (Fig. 3A; Fig. S8A). In all predicted complexes, the interaction surface was centered on short proline-rich motifs within the partner proteins (Fig. S8B). We therefore refer to this region of the TEsup proteins as a proline-recognition domain (PRD).

**Figure 3.**
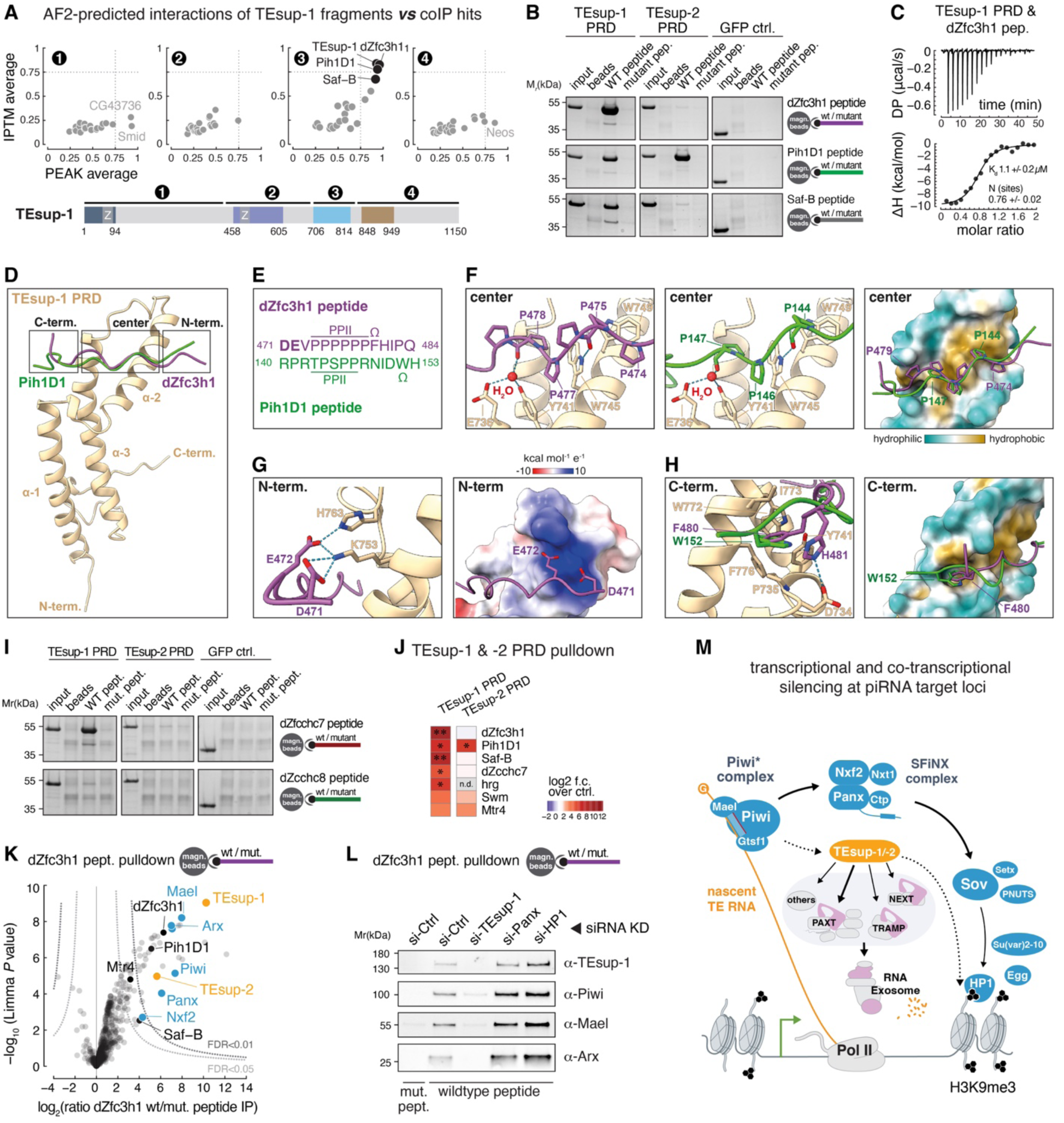
A polyproline recognition domain in TEsup-1 interacts with exosome adaptors. **(A)** Results from an AlphaFold2 Multimer pairwise interaction screen between indicated TEsup-1 fragments (1–4) and significantly enriched TEsup-1&2 interactors (FDR < 0.05) identified by co-IPs. Direct interaction confidence is indicated by interface predicted Template Modeling (IPTM) and PEAK scores (scaled and inverted maximum score from the Predicted Aligned Error (PAE) graph in the inter-protein quadrants) (Hohmann *et al*, 2025). **(B)** Coomassie-stained SDS–PAGE of in vitro peptide pulldown assays using streptavidin-bound wild-type or mutant dZfc3h1, Pih1D1, or Saf-B peptides, and recombinant GFP-tagged TEsup-1 PRD, TEsup-2 PRD, or GFP alone as prey. **(C)** Isothermal titration calorimetry (ITC) measurement of the interaction between the TEsup-1 PRD and the dZfc3h1 peptide. Binding affinity (*Kd*) and stoichiometry (N) represent mean ± s.d. from three biological replicates. **(D)** Crystal structures of the TEsup-1 PRD (pale yellow) in complex with proline-rich peptides from dZfc3h1 (purple) and Pih1D1 (green). The two complexes are superimposed; only one TEsup-1 PRD molecule is shown for clarity. The interaction interface is divided into N-terminal, central, and C-terminal regions. **(E)** Sequence alignment of proline-rich peptides from dZfc3h1 and Pih1D1 that bind the TEsup-1 PRD. The negatively charged region in dZfc3h1 is in bold; aromatic residues shown in (H) are labeled (Ω); regions adopting a PPII helix are highlighted. **(F)** Left: Zoom-in view of the central region of the TEsup-1 PRD–dZfc3h1 complex showing key interacting residues (sticks). Middle: corresponding view for the TEsup-1 PRD–Pih1D1 complex. Right: surface representation of TEsup-1 PRD colored by hydrophobic potential (brown, hydrophobic; cyan, hydrophilic), with central peptide regions (purple, green) shown and prolines indicated as sticks. **(G)** Left: Zoom-in view of the N-terminal region of dZfc3h1 bound to TEsup-1 PRD. Right: surface representation of TEsup-1 PRD colored by electrostatic potential (red, negative; white, neutral; blue, positive). Negatively charged dZfc3h1 residues contributing to the interface are shown as sticks. **(H)** Left: Zoom-in view of the C-terminal regions of dZfc3h1 and Pih1D1 peptides bound to TEsup-1 PRD. Right: surface representation of TEsup-1 PRD colored by hydrophobic potential (brown–cyan). The C-terminal peptide regions are shown as cartoons with aromatic residues highlighted. **(I)** As in (B), using dZcchc7 and dZcchc8 peptides and corresponding mutant versions. **(J)** Heatmap showing enrichment of indicated proteins in TEsup-1 and TEsup-2 FLAG co-Ips ( Table S1). Values represent means of three biological replicates (*FDR < 0.05, **FDR < 0.01 based on limma; n.d., not detected). **(K)** Volcano plot showing the log_2_-fold enrichment and significance of proteins identified by quantitative mass spectrometry in dZfc3h1 peptide pulldowns versus mutant controls (n = three biological replicates; Table S1). piRNA pathway factors (blue), TEsup proteins (yellow) and selected other factors (black) are labeled. **(L)** Western blot showing specific interaction of TEsup-1 and Piwi* factors with wildtype dZfc3h1 peptide, loss of interaction upon TEsup-1 or Piwi depletion, and enhanced enrichment upon Panx or HP1 depletion. **(M)** Schematic model showing the two silencing branches downstream of nuclear Piwi-piRNA complexes, centered on SfiNX and Creator proteins, respectively. Arrows indicate physical connections (bold: direct protein-protein interaction; dashed: potential connection).

Biochemical assays confirmed that the PRDs of TEsup proteins directly bind proline-rich motifs. Recombinant PRDs interacted with synthetic peptides derived from dZfc3h1, Saf-B, and Pih1D1 in vitro, whereas binding was abolished by targeted point mutations within the proline-rich sequences (Fig. 3B; Fig. S8C). Quantitative binding measurements revealed distinct affinities: TEsup-1 bound the dZfc3h1 peptide with low-micromolar affinity, whereas its interaction with Pih1D1 was considerably weaker (Fig. 3C; Fig. S8D).

To define the structural basis of these interactions, we determined crystal structures of the TEsup-1 PRD bound to peptides from dZfc3h1 (1.9 Å resolution) or Pih1D1 (2.1–2.3 Å resolution) (Fig. 3D–H; Fig. S8E; Data Table 1). The PRD adopts a compact tri-helical fold that presents a contiguous hydrophobic surface enriched in aromatic residues (Fig. 3D). In both complexes, the bound peptides adopt a left-handed, all-trans polyproline II (PPII) helical conformation (Fig. 3E,F). The dZfc3h1 peptide engages the PRD through extensive hydrophobic and aromatic contacts, including insertion of a C-terminal aromatic residue into a defined hydrophobic pocket (Fig. 3F,G). By contrast, the Pih1D1 peptide lacks additional N-terminal contacts, providing a structural explanation for its reduced binding affinity (Fig. 3F,H).

Consistent with a broader capacity for exosome adaptor engagement, the TRAMP and NEXT scaffolds dZcchc7 and dZcchc8 also harbor proline-rich motifs that bound robustly (dZcchc7) or moderately (dZcchc8) to the TEsup-1 PRD in vitro (Fig. 3I; Fig. S8F,G). Pull-down assays using recombinant PRDs and OSC lysates recapitulated these preferences, retrieving endogenous dZfc3h1, dZcchc7, Saf-B, and Pih1D1 (Fig. 3J). Moreover, co-immunoprecipitation experiments with dZcchc7 in cells depleted of the other adaptor scaffolds (dZfc3h1 and dZcchc8) resulted in enhanced recovery of both TEsup-1 and TEsup-2, supporting a model where adaptor scaffolds bind the same interface in TEsup proteins (Fig. S8H).

The biochemical and structural data suggested that TEsup proteins could bridge Piwi silencing complexes to the nuclear exosome via PRD-mediated adaptor binding. To test this possibility, we performed peptide pull-down experiments using dZfc3h1-derived peptides and nuclear OSC lysates (Fig. 3K). Strikingly, the wild-type peptide—but not a point-mutated version—retrieved TEsup-1 and TEsup-2, as well as all three Piwi* subunits (Piwi, Asterix, and Maelstrom) and the SFiNX components Panoramix and Nxf2 (Fig. 3K). Importantly, association of the dZfc3h1 peptide with the Piwi silencing machinery was lost upon depletion of TEsup-1 but increased when levels of target-engaged Piwi* were experimentally elevated by transposon de-silencing in Panoramix- or HP1-depleted cells (Fig. 3L; Fig. S8I).

To assess the functional relevance of the PRD *in vivo*, we generated in-frame deletions of the PRD at the endogenous *TEsup* loci in flies. Both mutant proteins were expressed at levels comparable to wild type and localized to the nucleus (Fig. S8J). While *TEsup-2^ΔPRD^* mutants displayed no detectable defects in TE silencing (consistent with the weak adaptor binding observed *in vitro*), *TEsup-1^ΔPRD^* mutants phenocopied *TEsup-1* null alleles, exhibiting upregulation of multiple transposons (Fig. S8K). Combined deletion of both PRDs resulted in even stronger TE derepression and near-complete sterility (Fig. S8K,L).

Together, these findings define the proline-recognition domain as a key molecular interface through which TEsup proteins, particularly TEsup-1, engage nuclear RNA exosome adaptors and other client proteins possessing proline-rich peptides. This interaction provides a mechanistic basis by which TEsup proteins can link Piwi–piRNA target recognition to exosome recruitment, enabling the co-transcriptional degradation of nascent transposon RNAs (Fig. 3M).

### The nuclear RNA exosome enforces Piwi-mediated silencing through RNA decay

To test whether the nuclear RNA exosome contributes functionally to Piwi-mediated transposon repression, we depleted core exosome subunits, the RNA helicase Mtr4, or adaptor scaffold proteins in OSCs and quantified RNA levels of the piRNA-repressed TEs *mdg1* and *gypsy* by RT–qPCR (Fig. 4A). Depletion of the catalytic subunits Rrp6 or Dis3, or of the core exosome components Rrp40 or Rrp41, caused strong derepression of both transposons (Fig. 4A; Fig. S9A). Among adaptor scaffolds, loss of dZfc3h1 increased *gypsy* RNA levels, whereas depletion of dZcchc7 or dZcchc8 individually had little effect (Fig. 4A; Fig. S9A,B). In contrast, co-depletion of any two adaptor scaffolds, all three scaffolds, or their shared helicase Mtr4 resulted in robust TE RNA accumulation comparable to depletion of core exosome subunits (Fig. 4A).

**Figure 4.**
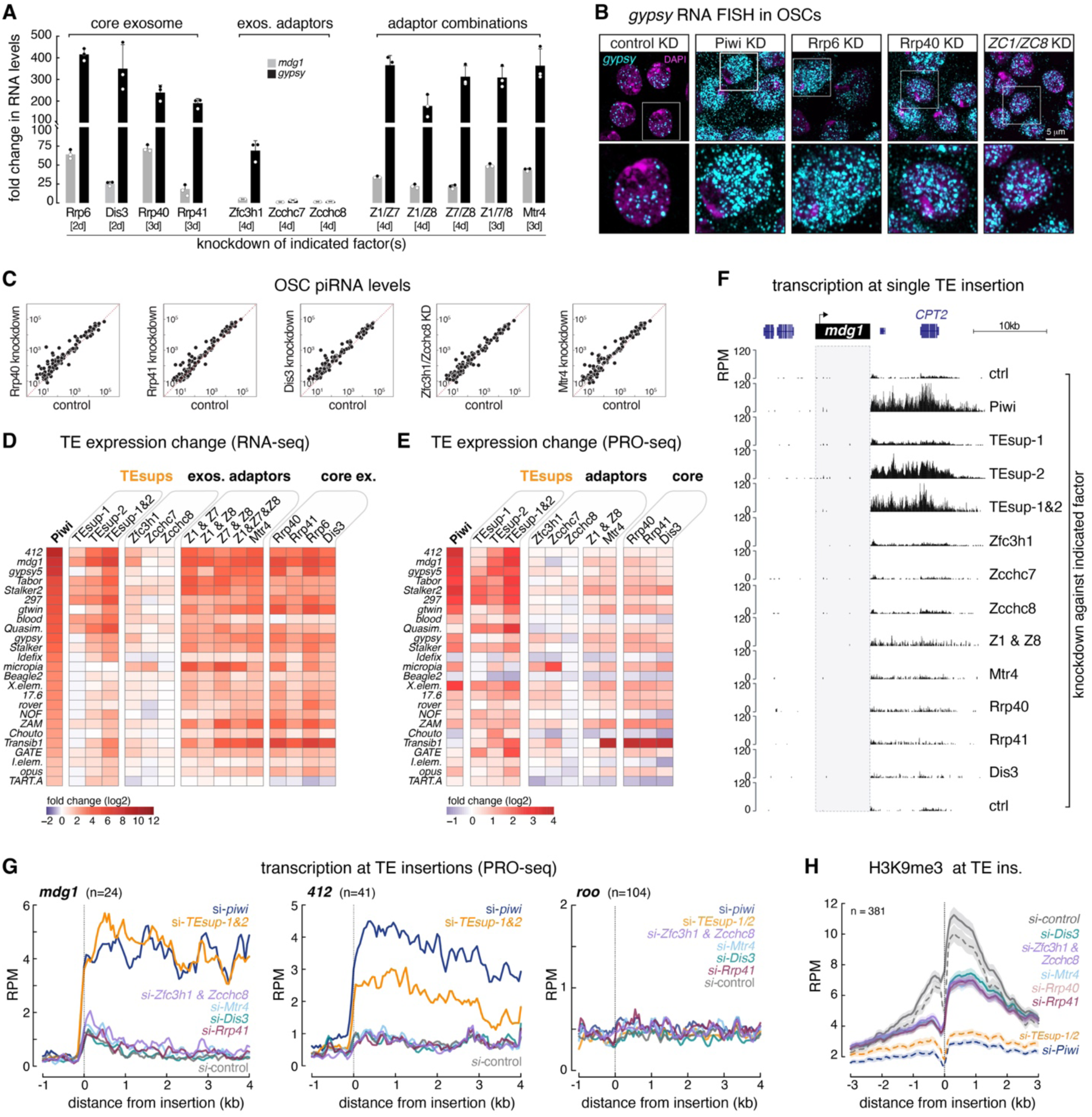
The nuclear RNA exosome is required for silencing piRNA-targeted transposons. **(A)** RT-qPCR analysis showing fold changes in steady-state RNA levels of indicated TEs in OSCs after siRNA-mediated depletion of nuclear RNA exosome-related factors (knockdown duration indicated in days). Data are normalized to si-*Luc* control and *act5C* mRNA levels (mean ± s.d., n = 3 biological replicates). **(B)** Confocal images of OSCs depleted for indicated factors by siRNAs and stained for DNA (DAPI) and *gypsy* TE RNA (FISH). Enlarged views of boxed regions are shown below. **(C)** Scatter plots showing antisense piRNA levels per TE in OSCs depleted for indicated factors versus control cells (small RNA-seq). **(D)** Heatmap showing log_2_-fold changes in steady-state RNA levels of Piwi-repressed TEs in OSCs after indicated knockdowns (ribo-zero RNA-seq, mean of n = 3 biological replicates). **(E)** Heatmap showing log_2_-fold changes in nascent transcriptional output of Piwi-repressed TEs in OSCs after indicated siRNA knockdowns (PRO-seq, mean of n = 2 biological replicates). **(F)** UCSC genome browser tracks showing nascent transcriptional activity (PRO-seq) at a euchromatic *mdg1* insertion after indicated knockdowns (n = 2 biological replicates; multi-mapping reads within the *mdg1* insertion excluded). **(G)** Metaplots of PRO-seq signal around *mdg1*, *412*, and *roo* insertions following depletion of indicated factors (insertions near endogenously expressed genes excluded). **(H)** Metaplot of H3K9me3 levels flanking piRNA-targeted TE insertions in OSCs after depletion of indicated nuclear RNA exosome-related factors (ChIP-seq, average of n = 2 biological replicates; n = 381 TE insertions; dashed lines indicate same dataset as in Fig. 2D).

To examine the cellular fate of TE transcripts upon exosome impairment, we performed single-molecule RNA FISH for *gypsy* in OSCs (Fig. 4B). In wildtype cells, we detected multiple bright nuclear foci, indicating substantial transcription of *gypsy* insertions in the presence of a functional piRNA pathway. However, no FISH signal could be detected in the cytoplasm, indicating rapid turnover of *gypsy* transcripts. Notably, depletion of exosome core subunits or of the adaptor complexes dZfc3h1 dZcchc8 together caused strong accumulation of *gypsy* transcripts in nucleus and cytoplasm (Fig. 4B; Fig. S9C), similar to the phenotype seen in Piwi-depleted cells (Fig. 4B; Fig. S9C). As depletion of exosome subunits, Mtr4, or adaptor scaffolds did not affect piRNA levels in OSCs (Fig. 4C), these results support a function of the exosome in Piwi-mediated silencing.

To assess exosome involvement globally, we performed RNA-seq in cells depleted of Piwi, TEsup proteins, or exosome components. Loss of core exosome subunits or Mtr4 caused strong upregulation of Piwi-repressed transposons (Fig. 4D; Fig. S9D-F). Individual adaptor knockdowns had minor effects, but co-depletion of two or all three scaffolds caused robust TE de-repression (Fig. 4D).

Piwi-mediated silencing is associated with transcriptional repression and heterochromatin formation (Le Thomas *et al*., 2013; Rozhkov *et al*., 2013; Sienski *et al*., 2012). Accordingly, depletion of Piwi resulted in elevated transcriptional output of its target TEs, as measured by PRO-seq (Fig. 4E) and Pol II ChIP-seq (Fig. S6B), including extended transcription downstream of individual TE insertions (Fig. 4E–G; Fig. S9G–L). Co-depletion of TEsup-1 and TEsup-2 produced an almost identical transcriptional phenotype (Fig. 4E–G; Fig. S9G–L), consistent with their established role in Piwi-dependent transcriptional silencing (Fig. 4H; Fig. S6A). In contrast, depletion of core exosome subunits, Mtr4, or adaptor scaffolds caused little or no increase in transcriptional activity at piRNA-targeted loci (Fig. 4E–G; Fig. S9G–N). Thus, the pronounced accumulation of Piwi-repressed TE RNAs upon exosome impairment is not due to increased transcription and is instead most consistent with defective nuclear RNA degradation.

Consistent with this interpretation, H3K9me3 levels at piRNA-repressed TEs were strongly reduced in Piwi- or TEsup-depleted cells but were only modestly affected by loss of exosome components or adaptor scaffolds (Fig. 4H). Together, these results establish the nuclear RNA exosome as a critical effector of Piwi-mediated transposon repression in OSCs. Whereas Piwi and TEsup proteins are also required for transcriptional silencing and heterochromatin formation, the exosome acts downstream to eliminate nascent TE transcripts, thereby enforcing silencing through co-transcriptional RNA decay.

### *P*-element silencing depends on Piwi–TEsup–exosome-mediated decay of nascent RNA

Having established a functional connection between Piwi, TEsup proteins, and the nuclear RNA exosome in cultured cells, we next examined whether this pathway operates *in vivo*. In ovaries, endogenously GFP-tagged scaffold proteins of the PAXT (dZfc3h1), TRAMP (dZcchc7), and NEXT (dZcchc8) complexes localized to nuclei, as observed in OSCs (Fig. S10A). Co-immunoprecipitation with tagged dZfc3h1 recovered PAXT components together with TEsup-1, TEsup-2, Piwi, and known Piwi cofactors, confirming that the physical associations identified in cultured cells are preserved in ovarian tissue (Fig. S10B).

To assess the functional relevance of the nuclear exosome for transposon silencing *in vivo*, we depleted adaptor scaffolds in the germline using validated shRNA transgenes driven by MTD-GAL4 (Fig. 5A; Fig. S10A,C). Individual knockdown of dZfc3h1, dZcchc7, or dZcchc8 caused little or no derepression of three well-characterized Piwi-repressed transposons (*blood*, *HeT-A*, and *mdg3*) (Fig. 5A). In contrast, combined depletion of dZfc3h1 with dZcchc7—and to a lesser extent with dZcchc8—resulted in strong TE upregulation (Fig. 5A), closely mirroring the cooperative requirement for adaptor complexes observed in OSCs. These results indicate that adaptor-mediated recruitment of the exosome is also essential for germline TE silencing *in vivo*.

**Figure 5.**
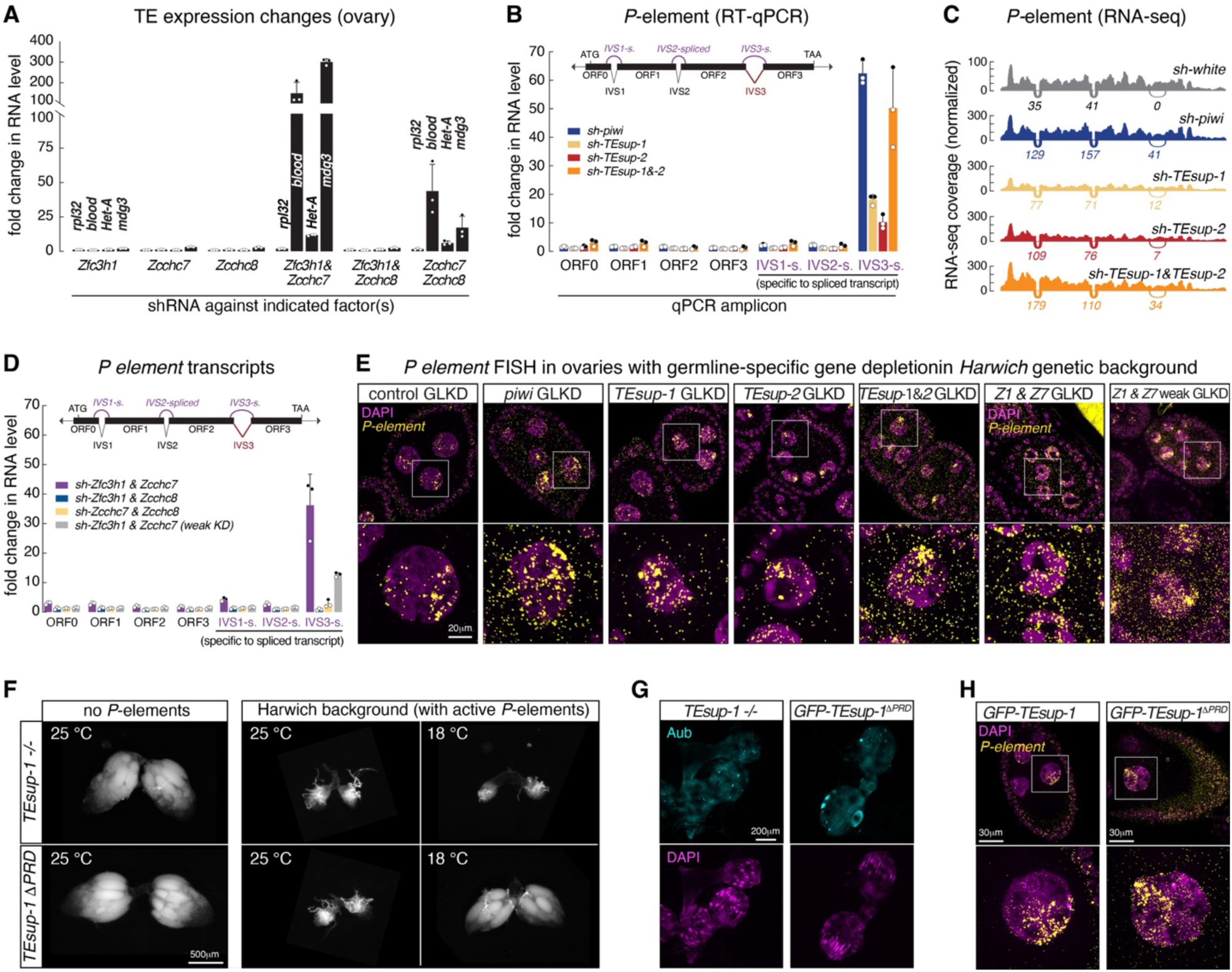
*P*-element silencing in the germline requires nascent RNA decay by the TEsup-exosome axis. **(A)** RT-qPCR analysis showing fold changes in steady-state RNA levels of indicated germline TEs in ovaries after depletion of indicated nuclear RNA exosome adaptor scaffolds using *MTD*-GAL4 driven shRNA transgenes. Data are normalized first to *rpl32* mRNA level and then to sh-*white* control (mean ± s.d., n = 3 biological replicates). **(B)** RT-qPCR analysis showing fold changes in *P*-element RNA levels using primers detecting the four coding exons (ORF0-3) and primers specific for spliced products of the three introns (IVS1-3) in ovaries depleted of indicated factors in the germline using *nanos*-GAL4 driven shRNA transgenes. Data are normalized first to *rpl32* mRNA level and then to sh-*white* control (mean ± s.d., n = 3 biological replicates). **(C)** Density plots of *P-element* RNA-seq reads (polyA+ libraries) (average normalized read counts of n = 3 biological replicates) in ovaries depleted of indicated factors in the germline using *nanos*-GAL4 driven shRNA transgenes. The number and position of spliced-reads across IVS1, IVS2 and IVS3 splicing junctions are shown below each plot. **(D)** As in (B), for ovaries depleted of indicated exosome adaptor scaffolds in the germline. **(E)** Confocal images of egg chambers from *Harwich* flies depleted for indicated factors in the germline using *nanos*-GAL4 driven shRNA transgenes, stained for DNA (DAPI) and *P*-element RNA (FISH). Enlarged views of boxed regions are shown below. **(F)** Bright-field images of ovaries from *TEsup-1* null (top row) or *TEsup-1^ΔPRD^* (bottom row) flies either lacking *P*-elements (left) or carrying *P*-elements from the *Harwich* background (right), reared at 25 °C or 18 °C (as null mutations were lethal, an equivalent analysis for *TEsup-2* was not possible). **(G)** Confocal images of rudimentary ovaries lacking germline tissue from indicated *TEsup-1* mutant flies carrying *P*-elements from the *Harwich* background reared at 25 °C, stained for Aubergine (top) and DNA (DAPI). **(H)** As in (B), for ovaries from *GFP-TEsup-1^ΔPRD^* flies (right) and the corresponding *GFP-TEsup-1* wildtype control, both carrying *P*-elements from the *Harwich* background and reared at 18 °C.

We next focused on an unresolved question in piRNA biology: how Piwi represses the *P*-element, a DNA transposon that largely escapes heterochromatin-based transcriptional silencing (Teixeira *et al*., 2017)? Unlike most transposons, the *P*-element lacks its own enhancer and instead depends on nearby host enhancers for transcriptional activation (O’Kane & Gehring, 1987), likely explaining its resistance to transcriptional repression. Previous work showed that loss of Piwi in the germline causes only modest increases in total *P*-element RNA but selectively enhances splicing of the third intron (IVS3), thereby enabling production of functional transposase (Teixeira *et al*., 2017). While these results suggested that Piwi suppresses *P*-element activity by blocking IVS3 splicing (Ghanim *et al*, 2020; Teixeira *et al*., 2017), the underlying molecular mechanism has remained unclear.

Depletion of TEsup proteins phenocopied Piwi loss, causing the same specific increase in IVS3 splicing (Fig. 5B,C). Strikingly, co-depletion of the exosome adaptor scaffolds dZfc3h1 and dZcchc7 resulted in an equivalent splicing phenotype (Fig. 5D; Fig. S10D). Given the established interaction between TEsup proteins and exosome adaptors, these results suggest that Piwi-dependent repression of the *P*-element does not occur through direct inhibition of splicing, but instead through rapid degradation of nuclear *P*-element transcripts prior to productive processing.

Single-molecule RNA FISH provided direct support for this model. In somatic follicle cells, where the *P-element* is not targeted by the piRNA pathway but IVS3 splicing is constitutively blocked by the somatic factor Psi (Adams *et al*, 1997), *P-element* transcripts accumulated in the cytoplasm, indicating that unspliced RNA is efficiently exported (Fig. S10E). In contrast, in germline cells, *P*-element transcripts were rarely detected in the cytoplasm despite prominent nuclear transcription foci, consistent with rapid nuclear turnover of nascent RNA (Fig. S10E). Loss of Piwi resulted in robust cytoplasmic accumulation of *P*-element transcripts (Fig. 5E).

Depletion of TEsup-1 or TEsup-2 produced an equivalent phenotype, and combined loss of both factors phenocopied Piwi depletion (Fig. 5E). Similarly, co-depletion of dZfc3h1 and dZcchc7 led to nuclear export of *P*-element transcripts (Fig. 5E; Fig. S10F,G). This phenotype was also observed using a weaker RNAi line, arguing against indirect effects arising from disrupted oogenesis (Fig. 5E; Fig. S10D). Together, these data indicate that Piwi suppresses *P*-element activity by promoting rapid co-transcriptional degradation of nascent *P*-element RNAs via the nuclear exosome.

Genetic evidence further supports this decay-centered mechanism. In the absence of *P*-elements, *TEsup-1* mutant flies developed ovaries of normal morphology, laid eggs, and were subfertile (Fig. S8L). In contrast, introgression of the *TEsup-1* null allele into flies carrying active *P*-elements uncovered a strong genetic interaction. Whereas wild-type flies harboring active but piRNA-repressed *P*-elements were fully fertile, *TEsup-1* mutants exhibited severe ovarian atrophy (Fig. 5F), germline loss (Fig. 5G), and complete sterility—hallmarks of P–M hybrid dysgenesis (O’Kane & Gehring, 1987; Teixeira *et al*., 2017). Strikingly, *P* cytotype flies carrying a GFP-tagged *TEsup-1^ΔPRD^* allele showed identical defects (Fig. 5F), whereas flies expressing a wild-type GFP–*TEsup-1* allele were fully fertile (Fig. S8L). At 18 °C, when *P*-element transposase activity is reduced (Ghanim *et al*., 2020), *TEsup-1^ΔPRD^* mutants remained sterile but developed ovaries with residual germline tissue (Fig. 5F). In these ovaries, nuclear export of *P*-element transcripts and accumulation of IVS3 spliced RNA were both elevated (Fig. 5H; Fig. S10H).

Together, these *in vivo* findings provide strong evidence that Piwi silences the *P*-element through TEsup-mediated recruitment of the nuclear RNA exosome, resulting in rapid co-transcriptional degradation of nascent transcripts. This decay-centered mechanism explains how Piwi can effectively repress a transposon that largely evades heterochromatin-based transcriptional silencing.

## DISCUSSION

Transposon silencing by nuclear Argonaute proteins presents a long-standing paradox: effective repression requires the elimination of transposon transcripts, yet Piwi must bind these same RNAs to maintain transcriptional repression and heterochromatin formation. Our results show that the *Drosophila* piRNA pathway resolves this conundrum by coordinating heterochromatin formation with co-transcriptional RNA degradation. This dual mechanism ensures that transposons are efficiently silenced even as they are transcribed.

Central to this dual control mechanism are the previously uncharacterized proteins TEsup-1 and TEsup-2, which play distinct yet complementary roles. On one hand, they promote transcriptional silencing and heterochromatin formation; on the other, they physically interact with nuclear RNA exosome adaptors. Our data indicate that TEsup-2 plays a dominant role in transcriptional silencing and heterochromatin establishment, whereas TEsup-1 primarily mediates exosome recruitment through a proline-recognition domain (PRD) that engages the adaptor scaffolds of the PAXT, TRAMP, and NEXT complexes.

Sequence and structural analyses indicate that TEsup proteins are restricted to Arthropoda, with one to four paralogs found across species. Notably, their proline-recognition domain (PRD) comprises a tandem repeat of a conserved short motif enriched in aromatic residues. A search of the Reference Proteome using the PRD HMM profile reveals significant similarity to the A/B boxes of FUBP proteins. FUBP proteins are widely conserved across eukaryotes and bind poly-proline motifs in partner proteins implicated in splice-site selection (Ebersberger *et al*, 2023).

Remarkably, the *Drosophila* FUBP homolog, Psi, functions as a splicing inhibitor of the *P*-element in somatic cells by engaging a proline motif in U1-70K via its C-terminal A/B boxes (Ignjatovic *et al*, 2005). The recruitment of RNA effector proteins via A/B box-like PRDs therefore seems to be a theme that cells repeatedly use for the co-transcriptional control of gene expression.

Loss of exosome activity in cultured cells causes a strong accumulation of Piwi regulated transposon RNAs but only modest changes in transcription activity, indicating that Piwi-mediated TE repression requires exosome-mediated RNA-decay, besides the well-established transcriptional silencing via local heterochromatin formation. Exosome depletion nonetheless produces a measurable reduction in H3K9me3 levels at piRNA target loci. Intact TE insertions are at the intersection of two opposing forces: Transcriptional activators promote their productive expression while the piRNA pathway seeks to suppress exactly this activity. We propose that rapid degradation of nascent, Piwi-bound transcripts is essential to maintain a repressive chromatin state: when RNA decay is impaired, nascent transcripts may accumulate and render TE chromatin more accessible to activating factors. This model is consistent with observations that the nuclear RNA exosome is indispensable for many chromatin-based silencing systems (Buhler *et al*, 2007; Khanduja *et al*, 2024; Zofall *et al*, 2012). It may also explain why undifferentiated cells, such as somatic stem cells or germline cells, are particularly sensitive to loss of TEsup proteins (Fig. 1G; Fig. S3C), as their chromatin is inherently more permissive to transcription to preserve developmental plasticity (Gaspar-Maia *et al*, 2011). Furthermore, the accumulation of non-degraded piRNA target transcripts may in turn lead to the futile sequestration of complementary Piwi-piRNA complexes or downstream silencing factors, ultimately resulting in global TE silencing defects.

All three exosome adaptor systems (PAXT, TRAMP, and NEXT) contribute to piRNA-mediated RNA decay, despite acting on largely distinct RNA classes in other contexts. This functional redundancy might reflect the diversity of transposon transcripts and their chromatin contexts, ensuring that even highly expressed or structurally atypical transposons are efficiently degraded.

The coupling of Piwi and TEsup proteins to the exosome also provides an explanation for how Piwi represses the *P*-element, a DNA transposon that largely evades heterochromatin-based silencing (Teixeira *et al*., 2017). Our results suggest that Piwi and TEsup proteins direct nascent *P*-element transcripts to the exosome for rapid degradation, thereby preventing accumulation of spliced, transposase-encoding mRNAs. In this case, Piwi does not block transcription directly but enforces silencing through accelerated RNA decay.

Although TEsup proteins physically associate with the Piwi silencing machinery, the mechanism underlying their recruitment to piRNA target loci remains unclear. Structural modeling did not reveal a high-confidence interface with Piwi or its cofactors. This might indicate that TEsup proteins promisciously associate with nascent transcripts and become stabilized or activated when Piwi engages the same RNA. In this convergent model, Piwi binding would act as a switch that diverts a nascent transcript from productive processing and export to decay, integrating small-RNA–guided recognition with RNA fate determination.

More broadly, our findings establish co-transcriptional RNA degradation as a key arm of nuclear Piwi function and reveal that the piRNA pathway enforces genome defense through integrated transcriptional and RNA-level repression. This principle likely reflects an ancient logic: the nuclear exosome contributes to heterochromatin-mediated silencing from fission yeast to mammals, and analogous coupling between chromatin repression and RNA decay occurs in the mammalian PRC2 and HUSH complexes (Buhler *et al*., 2007; Egan *et al*, 2014; Garland *et al*, 2022; Khanduja *et al*., 2024; Lee *et al*, 2013; Wendte *et al*, 2017; Zhang *et al*, 2014; Zhang *et al*, 2011; Zofall *et al*., 2012).

In summary, our study reveals the Piwi–TEsup–exosome axis as a critical silencing axis within the nuclear Piwi–piRNA pathway. By coupling transcriptional repression to RNA degradation, Piwi integrates chromatin regulation with RNA quality control, safeguarding genome integrity against the persistent threat of transposons.

## MATERIALS AND METHODS

### Fly strains and maintenance

All *Drosophila melanogaster* strains used in this study are listed (Table S2). The newly generated strains are available through VDRC (http://stockcenter.vdrc.at/control/main). All fly strains were maintained at 25 °C under a 12-hour light/dark cycle, except *P-element* related crosses as indicated at 18 °C. For fertility scoring and ovary analysis, flies were aged for 2-3 days and placed on apple juice agar supplemented with fresh yeast paste for 2 days.

N-terminal endogenous tagging of *CG31510/TEsup-1*, *CG7065/TEsup-2, CG4294/dZfc3h1* and C-terminal endogenous tagging of *dZcchc7*, *dZcchc8* with a 3xFLAG-V5-GFP tag were achieved through co-injection of a repair template and the pDCC6b plasmid (Addgene 59985) into *w^1118^* flies. The pDCC6b plasmid encodes both the guide RNA (gRNA) and Cas9 protein necessary for the CRISPR/Cas9-mediated integration. Guide RNAs used are listed (Table S3). Germline specific gene knockdowns were achieved by crossing shRNA transgene strains with the MTD-GAL4 driver line.

### OSC culture and transfection

Ovarian somatic cells (OSCs) were cultured in M3 medium supplemented with 10% fetal bovine serum, insulin, glutathione, and fly extract in a humidified incubator at 27°C with 2.5% CO_2_, following previously established protocols (Niki *et al*, 2006; Saito *et al*., 2009). The stable OSC reporter cell line, which harbors ten intronic boxB sites and 14 upstream UAS sites, was described in (Batki *et al*., 2019). Plasmid DNA and siRNA transfections were conducted using the MaxCyte electroporation system with the program "opt5". For small-scale transfections, 200 pmol of siRNA duplexes or 5 µg of plasmid DNA were used per 5 million cells. For larger scale transfections, 750 pmol of siRNA duplexes were used per 30 million cells. The siRNA duplexes used are listed (Table S4).

### Generation of endogenously tagged OSC cell lines

N-terminal endogenous tagging of Piwi, Panx, CG31510, and CG7065 with 3xFLAG-TurboID was performed by co-transfecting wild-type OSCs with Cas9 protein, the respective guide RNAs, and homologous directed repair (HDR) cassettes. These cassettes consisted of a puromycin-resistance gene followed by a P2A cleavage site linked to the 3xFLAG-TurboID biotin ligase.

For C-terminal tagging, TurboID-3xFLAG was inserted into Mael by co-transfecting WT OSCs with Cas9 protein, the appropriate guide RNA, and an HDR cassette encoding TurboID-3xFLAG followed by a P2A cleavage site linked to a puromycin-resistance gene. The OSC line expressing a nuclear 3xFLAG-TurboID-GFP fusion protein at comparable levels was generated to serve as a control for the TurboID experiments. For the N-terminal tagging of Panx, CG31510, and CG7065 with lN-3xFLAG, HDR cassettes containing a puromycin-resistance gene followed by a P2A cleavage site linked to λN-3xFLAG were introduced into the OSC reporter cell line. Similarly, N-terminal tagging of CG31510, CG7065, and CG4294/dZfc3h1 with 3xFLAG-V5-GFP was achieved by transfecting WT OSCs with HDR cassettes comprising a puromycin-resistance gene followed by a P2A cleavage site linked to 3xFLAG-V5-GFP. For C-terminal tagging, GFP-V5-3xFLAG was used for dZcchc7 and CG4622/dZcchc8, and dTAG-V5-3xFLAG was used for Mtr4. The respective HDR cassettes consisted of GFP-V5-3xFLAG or dTAG-V5-3xFLAG followed by a P2A cleavage site linked to a puromycin-resistance gene. These HDR cassettes were flanked by approximately 70 bp homology arms around the start or stop codon of the target genes. For each transfection, 5 µg of Cas9 protein, 12 µg of the relevant guide RNAs (Table S3), and 2.5 µg of the relevant purified HDR PCR product were used per 5 million OSCs. Two days post-transfection, the cells were split and subjected to selection with puromycin (5 µg/mL).

Puromycin-resistant clones were allowed to grow for 6-10 days, after which individual clones were picked and expanded. Individual clones were genotyped by PCR and western blot for successful and correct integration of the tag. OSC cell-lines generated in this study are listed (Table S2).

### Reporter tethering assay

The reporter tethering assay was performed in the OSC reporter cell line as previously described (Batki *et al*., 2019). Briefly, the coding sequences (CDS) of the proteins of interest were cloned into vectors (Addgene 128011 and 128014) to generate N-terminally λN- or Gal4-tagged fusion proteins. These λN/Gal4 fusion constructs were then transfected into OSC reporter cells at a concentration of 5 µg per 5 million cells, with the empty λN/Gal4 vector serving as a negative control. Four days post-transfection, the cells were harvested and analyzed by flow cytometry using a FACS BD LSR Fortessa (BD Biosciences). GFP fluorescence intensity was measured in the transfected cells, which were gated based on mCherry expression. Data analysis was conducted using FlowJo (FlowJo, LLC). Fold changes in GFP intensity, compared to control experiments, were calculated from 10,000 cells per experiment and visualized using RStudio (RStudio-2024.04.2-764).

### Generation of TEsup-1 and TEsup-2 specific monoclonal antibodies

His-tagged recombinant proteins of TEsup-1 (aa 950-1150) and TEsup-2 (aa 159-595) were used to generate mouse monoclonal antibodies specific to each protein. For TEsup-1, a single monoclonal antibody was produced for use in western blotting. For TEsup-2, one monoclonal antibody was generated for western blotting, and an additional monoclonal antibody was produced for immunofluorescence applications. All antibodies were raised at the MFPL Monoclonal Antibody Facility.

### Nuclei isolation and cell lysate preparation

OSCs were harvested following trypsinization and washed once with cold 1× PBS. For nuclei isolation, the cells were resuspended in swelling buffer (10 mM Tris-HCl, pH 7.5; 2 mM MgCl_2_; 3 mM CaCl_2_; Complete Protease Inhibitor Cocktail, Roche) and incubated on ice for 10 minutes. The cells were then centrifuged at 500 × g for 5 minutes at 4°C. The resulting cell pellet was resuspended in a gentle nuclei isolation buffer (10 mM Tris-HCl, pH 7.5; 2 mM MgCl_2_; 3 mM CaCl_2_; 0.25% IGEPAL CA-630; 10% glycerol; Complete Protease Inhibitor Cocktail, Roche) for co-immunoprecipitation and TurboID experiments. Nuclei were pelleted by centrifugation at 700 × g for 5 minutes at 4°C. For nuclear lysate preparation, the isolated nuclei were resuspended in an IP lysis buffer compatible with co-immunoprecipitation (50 mM HEPES, pH 7.3; 150 mM KCl; 3 mM MgCl_2_; 0.1% Sodium Deoxycholate; 0.25% Triton X-100; 0.25% IGEPAL CA-630; 10% glycerol; Benzonase; Complete Protease Inhibitor Cocktail, Roche) or in ice-cold whole cell extract (WCE) buffer (50 mM Tris-HCl, pH 7.5; 150 mM NaCl; 0.1% SDS; 0.5% Sodium Deoxycholate; 1% Triton X-100; Benzonase; Complete Protease Inhibitor Cocktail, Roche; 25 nM NEM when relevant) for TurboID experiments. After a 1-hour incubation on ice, the samples were centrifuged at 18,000 × g for 10 minutes at 4°C, and the nuclear lysate (supernatant) was collected. Protein concentration was measured using the Bradford assay. For whole cell extract preparation, cells were washed with cold PBS and directly resuspended in ice-cold WCE buffer. The suspension was incubated on ice for 30 minutes, followed by centrifugation at 18,000 × g for 10 minutes at 4°C. The WCE (supernatant) was collected, and protein concentration was measured by the Bradford assay.

### Western blot

15 µg of denatured proteins were separated by SDS-PAGE and transferred onto a 0.45 µm nitrocellulose membrane (Amersham™ Protran®). The membrane was blocked for 1 hour in 5% non-fat milk prepared in PBX (0.05% Triton X-100 in 1× PBS), followed by an overnight incubation at 4°C with the relevant primary antibody. After three washes in PBX (10 minutes each), the membrane was incubated with an HRP-conjugated secondary antibody in 5% non-fat milk in PBX for 1 hour at room temperature. This was followed by three additional PBX washes (5 minutes each). The membrane was then incubated with Clarity Western ECL Blotting Substrate (Bio-Rad) for 5 minutes and imaged using the Bio-Rad ChemiDoc MP imaging system. The antibodies used are listed (Table S5).

### TurboID of endogenously tagged proteins

TurboID experiments were conducted following a protocol previously described,(Branon *et al*., 2018) with specific modifications. OSCs with endogenously FLAG-TurboID tags were incubated with 500 µM biotin (Sigma, B4501) for 2 hours prior to harvesting. Nuclear lysates, as described in the "Nuclei isolation and cell lysate preparation" section, were used for affinity purification with Streptavidin Magnetic Beads (Pierce™ #88817). The lysates were incubated with the beads overnight (16 hours) at 4°C. The beads were then washed as follows: 1) A 10-minute wash with WCE buffer (50 mM Tris-HCl, pH 7.5; 150 mM NaCl; 0.1% SDS; 0.5% Sodium Deoxycholate; 1% Triton X-100; Benzonase; Complete Protease Inhibitor Cocktail (Roche); 25 nM NEM when relevant) at 4°C. 2) An 8-minute wash with 2% SDS at room temperature. 3) An 8-minute wash with wash buffer 1 (50 mM HEPES, pH 7.3; 500 mM NaCl; 1 mM EDTA; 0.1% Sodium Deoxycholate; 1% Triton X-100). 4) An 8-minute wash with wash buffer 2 (10 mM Tris-HCl, pH 7.5; 250 mM LiCl; 1 mM EDTA; 0.5% Sodium Deoxycholate; 1% IGEPAL CA-630). 5) Five washes (6 minutes each) with non-detergent wash buffer (20 mM Tris-HCl, pH 7.5; 137 mM NaCl). After washing, 20% of the beads were boiled in 2× SDS-PAGE loading buffer for 6 minutes at 95°C for subsequent western blot analysis, while the remaining 80% of the beads were reserved for mass spectrometry analysis.

### Co-immunoprecipitation of endogenously tagged proteins

Nuclear lysates from OSCs expressing different endogenously tagged proteins, prepared as described in the "Nuclei isolation and cell lysate preparation" section, were used for co-immunoprecipitation. The lysates were incubated with anti-Flag M2 Magnetic Beads (Sigma, M8823) for 3 hours at 4°C. Following incubation, the beads were washed four times (10 minutes each) with IP lysis buffer (50 mM HEPES, pH 7.3; 150 mM KCl; 3 mM MgCl_2_; 0.1% Sodium Deoxycholate; 0.25% Triton X-100; 0.25% IGEPAL CA-630; 10% glycerol; Benzonase; Complete Protease Inhibitor Cocktail, Roche) and five times (6 minutes each) with non-detergent wash buffer (20 mM Tris-HCl, pH 7.5; 137 mM NaCl) at 4°C. After washing, 30% of the beads were eluted with 3×FLAG peptide or boiled in 2× SDS-PAGE loading buffer for 6 minutes at 95°C for subsequent western blot analysis, while the remaining 70% were reserved for mass spectrometry analysis.

### Mass spectrometry analysis

Beads from TurboID or co-immunoprecipitation experiments were resuspended in 50 µl of 100 mM ammonium bicarbonate (ABC), supplemented with 400 ng of lysyl endopeptidase (Lys-C, Fujifilm Wako Pure Chemical Corporation), and incubated for 4 hours on a Thermo-shaker at 1200 rpm at 37°C. The supernatant was then transferred to a fresh tube and reduced with 0.5 mM Tris 2-carboxyethyl phosphine hydrochloride (TCEP, Sigma) for 30 minutes at 60°C, followed by alkylation with 3 mM methyl methanethiosulfonate (MMTS, Fluka) for 30 minutes at room temperature, protected from light. The sample was subsequently digested overnight at 37°C with 400 ng trypsin (Trypsin Gold, Promega). Finally, the digest was acidified by adding trifluoroacetic acid (TFA, Pierce) to a final concentration of 1%. An aliquot of each sample was then analyzed by LC-MS/MS.

Peptides were analyzed using a nano HPLC system (UltiMate 3000 RSLC nano system or Vanquish Neo UHPLC-System) coupled to an Orbitrap Exploris 480 mass spectrometer, equipped with a FAIMS Pro interface and a Nanospray Flex ion source (all parts Thermo Fisher Scientific). Peptides were initially loaded onto a trap column (Thermo Fisher Scientific, PepMap C18, 5 mm × 300 μm ID, 5 μm particles, 100 Å pore size) using 0.1% TFA as the mobile phase. After loading, the trap column was switched in line with the analytical column (Thermo Fisher Scientific, PepMap C18, 75 μm ID × 50 cm, 2 μm, 100 Å). Peptides were eluted at a flow rate of 230 nl/min, starting with mobile phases of 98% A (0.1% formic acid in water) and 2% B (80% acetonitrile, 0.1% formic acid). The gradient was linearly increased to 35% B over the next 120 minutes, followed by a steep gradient to 95% B over 5 minutes. This was maintained for 5 minutes before ramping down over 2 minutes to the starting conditions of 98% A and 2% B for equilibration at 30°C.

The Orbitrap Exploris 480 mass spectrometer was operated in data-dependent mode, performing a full scan (m/z range 350-1,200, resolution 60,000, normalized AGC target 100% or 300%) at three different compensation voltages (CVs: -45V, -60V, and -75V). This was followed by MS/MS scans of the most abundant ions, with a cycle time of 0.9 seconds for CVs -45V and - 60V, and 0.7 or 0.9 seconds for CV -75V. MS/MS spectra were acquired using an isolation width of 1.0 or 1.2 m/z, a normalized AGC target of 200%, a minimum intensity set to 25,000, HCD collision energy of 30, a maximum injection time of 100 ms, and a resolution of 30,000. Precursor ions selected for fragmentation (including charge states 2-6) were excluded for 45 seconds. The monoisotopic precursor selection (MIPS) mode was set to peptide, and the "exclude isotopes" feature was enabled.

For peptide identification, the RAW files were processed using Proteome Discoverer (version 2.5.0.400, Thermo Scientific). All MS/MS spectra were searched using MSAmanda v2.0.0.19924 (Dorfer *et al*, 2014). The peptide mass tolerance and fragment mass tolerance were set to ±10 ppm, using tryptic enzymatic specificity without proline restriction, allowing for up to two missed cleavages. Peptide and protein identification was conducted in two steps. 1) Initial Search: The RAW files were initially searched against the *Drosophila melanogaster* database (dmel-all-translation-r6.50; 22,280 sequences; 20,378,800 residues), supplemented with common contaminants and sequences of tagged proteins of interest. Beta-methylthiolation on cysteine was used as a fixed modification. The results were filtered to achieve a 1% false discovery rate (FDR) at the protein level using the Percolator algorithm (Kall *et al*, 2007) integrated within Proteome Discoverer. A sub-database of proteins identified in this search was generated for further analysis. 2) Secondary Search: The RAW files were then searched against the created sub-database using the same parameters as the initial search, but also considering the following additional variable modifications: oxidation on methionine, deamidation on asparagine and glutamine, phosphorylation on serine, threonine, and tyrosine, glutamine to pyro-glutamate conversion at peptide N-terminal glutamine, and acetylation on the protein N-terminus. The localization of post-translational modification sites within the peptides was determined using the ptmRS tool, which is based on the phosphoRS tool (Taus *et al*, 2011). Identifications were again filtered to 1% FDR at both protein and peptide spectrum match (PSM) levels, with an additional Amanda score cut-off of at least 150.

Protein areas were computed using IMP-apQuant (Doblmann *et al*, 2019) by summing up unique and razor peptides. The resulting protein areas were normalized using intensity-based absolute quantification (iBAQ) (Schwanhausser *et al*, 2011), and either sum normalization was applied across samples or the normalization with multiplication by a factor per sample to obtain a median protein ratio of 1 between sample pairs. Match-between-runs (MBR) was applied for peptides with high-confidence peak areas that were identified by MS/MS spectra in at least one run.

Proteins were required to be identified by a minimum of 2 PSMs in at least 1 sample, and quantified proteins were filtered to include at least 3 quantified peptide groups.

The statistical significance of differentially expressed proteins was determined using the limma package (Smyth, 2004). Results were visualized using volcano plots, scatter plots, or heatmaps, plotted with RStudio (RStudio-2024.04.2-764). All mass spectrometry datasets and their ProteomeXchange/ PRIDE dataset identifiers are listed (Table S1).

### RNA preparation and RT-qPCR

Total RNA was isolated from 5–10 million OSCs or 5–10 pairs of ovaries using the NucleoSpin® RNA kit (Macherey-Nagel) following the manufacturer’s instructions. Complementary DNA (cDNA) was synthesized from 1 µg of total RNA using random hexamer primers and SuperScript IV Reverse Transcriptase (Invitrogen). The primers used for qPCR analysis of genes and TEs are listed (Table S6).

### RNA-seq with polyA selection

For RNA sequencing, 1 µg of total RNA was used for poly(A)+ RNA selection. The poly(A)+ RNA selection, cDNA synthesis, and library preparation were performed using the NEBNext® Poly(A) mRNA Magnetic Isolation Module (NEB, E7490) following the manufacturer’s instructions. The libraries were sequenced on an Illumina HiSeq V4/2500 or NovaSeqX platform.

### RNA-seq with rRNA depletion

rRNA depletion was performed following the method described by (Morlan *et al*, 2012), with modifications. Total RNA isolated using the NucleoSpin® RNA kit was further purified using the RNA Clean & Concentrator-5 kit with on-column DNase I digestion (Zymo Research) according to the manufacturer’s instructions. For the depletion process, 1 µg of purified total RNA was mixed with 4 µg of an antisense oligos mix targeting *Drosophila melanogaster* rRNAs (listed in Table S7) in RNase H Buffer (20 mM Tris-HCl, pH 8; 100 mM NaCl; 50 µM EDTA). The mixture was denatured at 95°C for 3 minutes, followed by annealing with a temperature gradient from 95°C to 45°C (0.1°C/second). Subsequently, the RNA-DNA hybrids were digested with RNase H (Epicentre) for 1 hour at 45°C. Excess antisense oligos were then digested with TURBO DNase (Invitrogen), and the rRNA-depleted RNA was purified using the RNA Clean & Concentrator-5 kit (Zymo). Libraries were prepared using the NEBNext® Ultra™ II Directional RNA Library Prep Kit for Illumina® (NEB) according to the user protocol and sequenced on a NextSeq 2000 platform (Illumina) in single-read 50 mode.

### RNA-seq analysis

Sequencing reads were de-multiplexed and trimmed to remove adaptor sequences. Reads mapping to the *Drosophila melanogaster* rRNA precursor and mitochondrial genome were excluded using Bowtie (v1.3.1) with zero mismatches. The remaining reads were mapped to the *Drosophila melanogaster* genome (dm6) using STAR v2.7.10b (Dobin *et al*, 2013) with specific settings optimized for accurate mapping (settings: --outSAMmode NoQS --readFilesCommand cat --alignEndsType Local --twopassMode Basic --outReadsUnmapped Fastx --outMultimapperOrder Random --outSAMtype SAM --outFilterMultimapNmax 1000 --winAnchorMultimapNmax 2000 --outFilterMismatchNmax 0 --seedSearchStartLmax 30 --alignSoftClipAtReferenceEnds No --outFilterType BySJout --alignSJoverhangMin 15 --alignSJDBoverhangMin 1). Genome-mapped reads were intersected with Flybase genome annotations (r6.40) (Ozturk-Colak *et al*, 2024) using Bedtools v2.28 (Quinlan & Hall, 2010).

Reads mapped to multiple genomic locations were assigned to annotations based on the highest count. Subsequently, reads mapped to rRNA, tRNA, small nuclear RNA, small nucleolar RNA loci, and the mitochondrial genome were excluded from further analysis.

For gene quantification and differential gene expression (DGE) analysis, filtered and unmapped reads were randomized for quantification using Salmon (v1.5.1) with appropriate bias correction settings (settings: --dumpEqWeights --seqBias -- gcBias --useVBOpt --numBootstraps 100 -l SF --incompatPrior 0.0 --validateMappings). Quantification was performed against an index containing Flybase genome annotations (r6.40) and TE-consensus sequences in both orientations. Gene length-corrected trimmed mean of M-values (GeTMM) values were generated using edgeR (v3.34.0). DGE analysis was performed pairwise between libraries using DESeq2 (v1.32.0). For the P-element splicing analysis, RNA-seq reads were aligned to the transposon consensus sequences with known P-element IVS locations supplied as annotated splice junctions in the STAR index. Data visualization, including volcano plots, scatter plots, and heatmaps, was conducted using RStudio (RStudio-2024.04.2-764). All sequenced libraries with their GEO accession numbers are listed (Table S8).

### Transient transcriptome sequencing (TT-seq)

The TT-seq experiment was performed following the protocol in (Schwalb *et al*, 2016) with modifications. Briefly, OSCs were cultured on plates after siRNA transfections. Before harvesting the siRNA-treated cells, they were incubated with 4mM nucleoside analog 4-thiouridine (4sU) for 1 hour. Total RNA was isolated with Trizol and fragmentated to 200 to 500 nt long fragments. Fragmentated total RNA was incubated with freshly prepared MTS-biotin solution, 4sU-labelled RNA was purified with streptavidin beads via the µMACS kit. µMACS purified 4sU-labelled RNA was then subjected for clean-up and concentration with the Zymo RCC5 kit, followed by Turbo DNase digestion. 4sU-labelled RNA was then depleted for rRNA and libraries were prepared as described in “RNA-seq with rRNA depletion”. All sequenced libraries with their GEO accession numbers are listed (Table S8)

### Small RNA-seq

Small RNA libraries were prepared using the TraPR method (Grentzinger *et al*, 2020) with modifications. Briefly, Argonaute–small RNA complexes from OSC lysates were isolated using TraPR ion exchange spin columns. The small RNAs were then purified via acidic phenol-chloroform extraction. Following purification, the small RNAs were ligated with adaptors at both the 3’ and 5’ ends (Jayaprakash *et al*, 2011), followed by reverse transcription and PCR amplification. The resulting small RNA libraries were sequenced on a NovaSeq 6000 platform (Illumina) in single-read 100 mode.

### Small RNA-seq analysis

After de-multiplexing by barcodes and trimming to remove adaptor sequences, the small RNA reads were mapped to the *Drosophila melanogaster* genome (dm6) using Bowtie (v1.3.1) (Langmead, 2010) with no mismatches allowed. Genome-mapped reads were intersected with Flybase genome annotations (r6.40) using Bedtools v2.28 (Quinlan & Hall, 2010). Reads mapped to multiple genomic locations were assigned to annotations based on the highest count. Reads mapping to rRNA, tRNA, small nuclear RNA, small nucleolar RNA loci, and the mitochondrial genome were excluded before quantifying piRNAs mapped to individual TEs. For piRNA quantification, reads were mapped to TE consensus sequences using Bowtie with no mismatches allowed. Reads with multiple mappings were excluded, except for reads mapping twice within the same TE, where they were counted only once. Small RNA counts on TEs were normalized to 1 million miRNA reads and TE length to obtain RPKM-like quantifications for individual TEs.

Data visualization, including scatter plots and histograms, was conducted using RStudio (RStudio-2024.04.2-764). All sequenced small RNA libraries with their GEO accession numbers are listed (Table S8).

### Immunofluorescence staining of ovaries

Flies aged 2-3 days were placed on apple juice agar with fresh yeast paste for 2 days before ovary dissection. Ten ovary pairs were dissected into ice-cold 1× PBS and subsequently fixed in fixation buffer (4% formaldehyde and 0.3% Triton X-100 in 1× PBS) with rotation for 20 minutes at room temperature. After fixation, the ovaries were washed three times (10 minutes each) with PBX (0.3% Triton X-100 in 1× PBS), followed by a 30-minute blocking step in BBX (1% BSA and 0.3% Triton X-100 in 1× PBS) at room temperature. The blocked ovaries were incubated with the primary antibody diluted in BBX overnight at 4°C with rotation. Following three 10-minute washes in PBX, the ovary samples were incubated with a fluorophore-conjugated secondary antibody diluted in BBX overnight at 4°C with rotation in the dark. After a 10-minute wash in PBX, the ovaries were stained with 0.1 µg/mL DAPI in PBX for 10 minutes, followed by three additional washes. The ovaries were then mounted and imaged using a Zeiss LSM-980 Axio Imager confocal microscope, and the images were processed with FIJI/ImageJ. For endogenously GFP-tagged proteins, ovaries were directly subjected to DAPI staining after fixation, bypassing the antibody incubation steps. Antibodies used in this procedure are listed (Table S5).

### Single molecule RNA fluorescent in situ hybridization

*mdg1, Quasimodo, Het-A*, *blood* and *P*-element single molecule RNA–FISH on ovaries and *gypsy* RNA FISH on OSCs were performed using Stellaris probes (Biosearch Technologies) as described (Baumgartner *et al*, 2022). Ten ovary pairs were dissected into ice-cold 1× PBS and subsequently fixed in fixation buffer (4% formaldehyde and 0.3% Triton X-100 in 1× PBS) with rotation for 20 minutes at room temperature. Following fixation, the ovaries were washed three times (5 minutes each) with PBX (0.3% Triton X-100 in 1× PBS) and incubated in 500 µl of 70% ethanol overnight at 4°C. The ovaries were then rehydrated in wash buffer (10% formamide in 2× SSC) for 5 minutes. For hybridization, the ovaries were incubated in 50 µl of hybridization buffer (100 mg/ml dextran sulfate and 10% formamide in 2× SSC) with 0.5 µl of 25 µM Stellaris probes overnight at 37°C. After hybridization, the ovaries were quickly rinsed twice in wash buffer and then washed twice at 37°C for 30 minutes each. The samples were subsequently stained with 0.2 µg/mL DAPI in 2× SSC for 5 minutes, followed by two additional washes with 2× SSC for 5 minutes each. The stained ovaries were mounted using DAKO mounting medium (Agilent) and equilibrated in the dark at room temperature overnight. The mounted ovaries were imaged using a Zeiss LSM-980 Axio Imager confocal microscope. Images were processed with FIJI/ImageJ, and a Z-stack projection of five slices was generated. *mdg1, Quasimodo, Het-A*, *blood, gypsy* and *P*-element probe sequences used in this procedure are listed (Table S7).

### Precision Run-On Sequencing (PRO-seq)

PRO-seq was performed according to (Mahat *et al*, 2016) with modifications. Briefly, 20 million OSC cells and 1 million human HCT 116 spike-in cells were washed with ice-cold 1× PBS and resuspended in ice-cold permeabilization buffer (10 mM Tris-HCl, pH 7.5; 300 mM sucrose; 10 mM CaCl_2_; 5 mM MgCl_2_; 1 mM EGTA; 0.05% Tween-20; 0.1% IGEPAL CA-630; 0.5 mM DTT; Complete Protease Inhibitor Cocktail, Roche). After a 5-minute permeabilization on ice, the cells were spun down at 1000× g for 5 minutes at 4°C and washed twice by resuspending in ice-cold permeabilization buffer. Nuclei were resuspended in 200 µl storage buffer (10 mM Tris-HCl, pH 8; 25% glycerol; 5 mM MgCl_2_; 0.1 mM EDTA; 5 mM DTT), snap-frozen in liquid nitrogen, and stored at -70°C until use. Nuclear transcription run-on (NRO) was carried out by adding 100 µl of 2× NRO buffer (10 mM Tris-HCl, pH 8; 5 mM MgCl_2_; 1 mM DTT; 300 mM KCl; 0.25 mM ATP; 0.25 mM GTP; 0.05 mM Biotin-11-CTP; 0.05 mM Biotin-11-UTP; 0.8 U/µl murine RNase inhibitor; 1% sarkosyl) to 10 million nuclei in 100 µl storage buffer, followed by incubation at 30°C for 3 minutes. The reaction was terminated by adding 500 µl Trizol LS reagent (Invitrogen), followed by RNA extraction with chloroform and ethanol precipitation. The RNA was resuspended in 50 µl water and denatured at 65°C for 40 seconds, then subjected to base hydrolysis with 5 µl NaOH (1 M) for 15 minutes. Base hydrolysis was quenched with 25 µl Tris-HCl (1 M, pH 6.8), and RNA was purified through a Bio-Rad P30 column. Biotinylated nascent RNA was enriched using 50 µl M280 streptavidin beads for 30 minutes at room temperature, followed by two 5-minute washes with 500 µl high salt buffer (50 mM Tris-HCl, pH 7.5; 2 M NaCl; 0.5% Triton X-100), two washes with 500 µl binding buffer (10 mM Tris-HCl, pH 7.5; 300 mM NaCl; 0.1% Triton X-100), and one wash with 500 µl low salt buffer (5 mM Tris-HCl, pH 7.5; 0.1% Triton X-100). The biotinylated nascent RNA on streptavidin beads was then extracted with Trizol, purified through a direct-zol column (Zymo), and eluted with 5 µl diluted 3’ RNA linker (12.5 µM), followed by T4 ligation overnight at 16°C. Biotinylated nascent RNA was enriched again using 12.5 µl M280 streptavidin beads with the previously described washes and treated with Cap-Clip Pyrophosphatase (Biozym) for 1 hour at 37°C on the beads, followed by T4 polynucleotide kinase (NEB) treatment at 37°C for 1 hour. After repeating the same washes as for streptavidin bead enrichment, T4 ligation with a 5’ RNA linker (12.5 µM) was performed on the beads at room temperature for 4 hours. Following the washes as described before, RNA was extracted with Trizol, purified through a direct-zol column (Zymo), and eluted with 10.5 µl water. The purified RNA was used for reverse transcription with Superscript III Reverse Transcriptase (Thermo) according to the manufacturer’s instructions, followed by PCR amplification with the KAPA HiFi real-time library amplification kit (KAPA Biosystems). Libraries were purified with AMPure beads and sequenced on a NextSeq 550 platform.

### PRO-seq analysis

Sequencing reads were de-multiplexed by barcodes and trimmed to remove adaptor sequences. After fixing the start nucleotide and performing strand switching, reads were mapped to the *Drosophila melanogaster* genome (dm6) using Bowtie v1.3.1 (Langmead *et al*, 2009) with no mismatches allowed. Genome-mapped reads were intersected with Flybase genome annotations (r6.40) using Bedtools v2.28 (Quinlan & Hall, 2010). Reads mapped to multiple genomic locations were assigned to annotations based on the highest count. Subsequently, reads mapped to rRNA, tRNA, small nuclear RNA, small nucleolar RNA loci, and the mitochondrial genome were excluded from further analysis.

For gene quantification and DGE analysis, the filtered and unmapped reads were randomized for quantification using Salmon (v1.5.1; settings: --dumpEqWeights --seqBias --gcBias –useVBOpt --numBootstraps 100 -l SF --incompatPrior 0.0 --validateMappings) against an index containing Flybase gene annotations (CDS only, r6.40) and TE-consensus sequences (both orientations).

GeTMM were generated using edgeR (v3.34.0). Pairwise DGE analysis between libraries was performed using DeSeq2 (v1.32.0). Data visualization in heatmaps was conducted using RStudio (RStudio-2024.04.2-764), with replicates averaged for analyses. All sequenced libraries with their GEO accession numbers are listed (Table S8).

### ChIP-seq

ChIP was performed according to the protocol by (Lee *et al*, 2006) with modifications. OSCs were cultured on plates and crosslinked with 1% formaldehyde in 1× PBS for 10 minutes at room temperature. The reaction was quenched with glycine (final concentration of 125 mM) for 10 minutes, followed by two quick washes with ice-cold PBS. Chromatin was prepared using lysis buffer (50 mM Tris-HCl, pH 8.0; 2 mM EDTA; 0.5% IGEPAL CA-630; 10% Glycerol; Complete Protease Inhibitor Cocktail, Roche) and sonicated in sonication buffer (20 mM Tris-HCl, pH 8.0; 150 mM NaCl; 2 mM EDTA; 0.5% SDS; Complete Protease Inhibitor Cocktail, Roche) using a Covaris E220 for 6 minutes (Duty Factor 5.0, Peak Incident Power 140, Cycles per Burst 200). The samples were then centrifuged at 18,000 × g for 10 minutes at 4°C. The supernatant chromatin was snap-frozen in liquid nitrogen and stored at -70°C until use for ChIP. Protein G and A Dynabeads (mixed 1:1) were blocked in 1× TE buffer with 1 mg/ml denatured yeast tRNA (Sigma-Aldrich) and 1 mg/ml BSA (NEB) for 2 hours at 4°C, and then coupled with 5 µg of anti-H3K9me3 antibody (Active Motif, 39161) or Pol II antibody (Anti-RNA Polymerase II, CTD, cl8WG16) for 2 hours at 4°C. Sheared chromatin was diluted fivefold with ChIP dilution buffer (16.7 mM Tris-HCl, pH 8.0; 167 mM NaCl; 1.2 mM EDTA; 0.5% IGEPAL CA-630; Complete Protease Inhibitor Cocktail, Roche) and incubated with antibody-coupled beads overnight at 4°C. After incubation, the beads were washed with low salt buffer (20 mM Tris-HCl, pH 8.0; 150 mM NaCl; 2 mM EDTA; 0.5% IGEPAL CA-630; 0.1% SDS), high salt buffer (20 mM Tris-HCl, pH 8.0; 500 mM NaCl; 2 mM EDTA; 0.5% IGEPAL CA-630; 0.1% SDS), and LiCl wash buffer (10 mM Tris-HCl, pH 8.0; 250 mM LiCl; 1 mM EDTA; 0.5% IGEPAL CA-630; 0.5% sodium deoxycholate). The ChIP chromatin was then eluted with elution buffer (0.1 M NaHCO_3_, 1% SDS). Eluates and inputs were de-crosslinked overnight at 65°C, treated with RNase A and proteinase K, and then purified using the ChIP DNA Clean & Concentrator kit (Zymo) according to the user manual. ChIP-seq libraries were prepared using the NEBNext Ultra DNA Library Prep Kit (NEB) and sequenced on a NovaSeq X Plus (Illumina) in SR100 mode or a NextSeq 2000 (Illumina) in SR50 mode.

### ChIP-seq analysis

ChIP-seq analysis was conducted as described in (Baumgartner *et al*., 2022). Briefly, following adaptor removal, ChIP-seq reads with a minimum length of 18 nt were mapped to the *Drosophila melanogaster* genome (dm6) using Bowtie v1.3.1. For genome-wide analysis, reads were mapped with zero mismatches allowed. For TE-consensus analysis, reads were mapped allowing zero mismatches, with multi-mapping permitted only within a single TE. BigWig files were generated using Homer (Heinz *et al*, 2010) and UCSC BigWig tools.(Kent *et al*, 2010) Meta profiles were generated using ngs.plot(Shen *et al*, 2014) with bam files containing uniquely mapped reads, and replicates were averaged for analyses. Piwi-regulated TEs were identified according to (Sienski *et al*, 2015). All sequenced libraries with their GEO accession numbers are listed (Table S8).

### Protein Expression and Purification

The DNA corresponding to the TEsup-1 PRD (aa 702-820), either alone or fused to dZfc3h1 (aa 471-482) through a (GS)6 linker, was cloned into a modified pET expression vector to generate an N-terminal His10-MBP-3C fusion protein. These constructs were expressed in *Escherichia coli* BL21(DE3) grown at 37°C in Terrific Broth medium until the culture reached an optical density at 600 nm of 2–3. Subsequently, the temperature was reduced to 18°C, and protein expression was induced with 0.2 mM IPTG for 14-16 hours overnight. Cell pellets were resuspended in lysis buffer (25 mM Tris-HCl, pH 8.0; 500 mM NaCl; 50 mM NaH_2_PO_4_; 25 mM Imidazole; 10% (v/v) Glycerol; 5 mM 2-Mercaptoethanol) supplemented with 1 mM PMSF and 4 units/ml DNase I. Cells were lysed by sonication for 3× 3 minutes using a Sonifier W-450 D (Branson) with the following settings: 50% amplitude, 1-second sonication, 1-second pause.

TEsup-1 PRD and TEsup-1 PRD-dZfc3h1 fusion proteins were purified with three purification steps. 1) Ni-NTA chromatography, followed by dialysis and tag cleavage with His-tagged 3C protease overnight during dialysis in containing 25 mM Tris-HCl, pH 8.0; 150 mM NaCl; 10% (v/v) Glycerol; 5 mM 2-Mercaptoethanol. 2) reverse Ni-NTA chromatography for removal of the 3C protease and the His10-MBP tag. 3) Gel filtration using a HiLoad Superdex 75 16/600 pg column in gel filtration buffer (25 mM Tris-HCl, pH 8.0; 150 mM NaCl; 2 mM DTT). For peptide pulldown and recombinant protein pulldown assays, the DNA corresponding to the TEsup-1 PRD (aa 702-820) or TEsup-2 PRD (aa 254-387) was cloned into a modified pET expression vector to generate an N-terminal His10-FLAG-GFP-3C-V5 fusion protein. These proteins were purified using the following three-step process. 1) Ni-NTA chromatography purification though a 5 ml HisTrap HP column (Cytiva, #17524801). 2) Anion exchange chromatography purification though HiTrap Q anion exchange column (Cytiva, #17115401). 3) Gel filtration purification using a HiLoad Superdex 75 16/600 pg column.

### Peptide pulldown recombinant protein or nuclear extract

Peptide pulldown was performed according to (Andreev *et al*., 2022). In brief, wild-type (WT) and mutant peptides corresponding to dZfc3h1 (aa 466-484), Pih1D1 (aa 135-154), and Saf-B (aa 643-660) were synthesized with an aminohexanoate-linked N-terminal biotin moiety and a C-terminal amide blocking group. Excess peptides were coupled to 12 µl streptavidin magnetic beads (Pierce) in 200 µl pulldown buffer (25 mM HEPES; 70 mM NaCl; 5% Glycerol; 0.01% IGEPAL CA-630; Complete Protease Inhibitor Cocktail, Roche) for 3 hours at 4°C, followed by three 5-minute washes with pulldown buffer. For pulldown with recombinant protein, the pre-coupled beads-peptide complexes were incubated with 28 µg of recombinant protein for 2 hours at 4°C, followed by three 30-second washes with 300 µl pulldown buffer. The beads were then boiled in 20 µl 2× SDS–PAGE loading buffer at 95°C for 5 minutes, and the eluate was analyzed by SDS–PAGE followed by Coomassie staining. For pulldown with OSC nuclear extract, 50 µl of beads were used. The pre-coupled beads-peptide complexes were incubated with 2 mg of OSC nuclear extract in IP lysis buffer (50 mM HEPES, pH 7.3; 150 mM KCl; 3 mM MgCl_2_; 0.1% Sodium Deoxycholate; 0.25% Triton X-100; 0.25% IGEPAL CA-630; 10% Glycerol; Benzonase; Complete Protease Inhibitor Cocktail, Roche) for 4 hours at 4°C. This was followed by four 10-minute washes in IP lysis buffer and five 6-minute washes with non-detergent wash buffer (20 mM Tris-HCl, pH 7.5; 137 mM NaCl) at 4°C. After washing, 30% of the beads were boiled in 30 µl 2× SDS-PAGE loading buffer for 6 minutes at 95°C for western blot analysis, while the remaining 70% of the beads were used for mass spectrometry analysis.

### Recombinant protein pulldown with OSC nuclear extract

Forty micrograms of recombinant His10-FLAG-GFP-3C-V5 fused TEsup-1 PRD (aa 702-820) or TEsup-2 PRD (aa 254-387) was coupled to 25 µl of GFP-Trap Magnetic Agarose (Chromotek) in IP lysis buffer at 4°C for 4 hours, followed by three 5-minute washes with IP lysis buffer. The pre-coupled beads-recombinant protein complexes were then incubated with 2 mg of OSC nuclear extract in IP lysis buffer for 4 hours at 4°C. This was followed by four 10-minute washes with IP lysis buffer and five 6-minute washes with non-detergent wash buffer at 4°C. After washing, 30% of the beads were boiled in 30 µl of 2× SDS-PAGE loading buffer for 6 minutes at 95°C for western blot analysis, while the remaining 70% of the beads were reserved for mass spectrometry analysis.

### Crystallization

Crystallization trials with the TEsup-1 PRD-dZfc3h1 fusion protein stored in gel filtration buffer (25 mM Tris-HCl pH 8.0; 150 mM NaCl; 2 mM DTT) were performed using a sitting-drop vapor diffusion set-up by mixing the sample and crystallization solution in a 1:1 and 2:1 ratio. The best diffracting crystals were obtained at a protein concentration of 9 mg/ml after seeding in condition G10 (Sodium acetate pH 4.6 and 1.1 M di-Ammonium tartrate) from the SaltRX screen (Hampton Research). For crystallization of the TEsup-1 RD/Pih1D1 complex, TEsup-1 and the Pih1D1 peptide were mixed in gel filtration buffer (25 mM Tris-HCl pH 8.0; 150 mM NaCl; 2 mM DTT), where Pih1D1 was added in 1.2 molar excess. Crystallization trials were performed as described above. The best diffracting crystals were obtained at a protein concentration of 14 mg/ml in condition D10 (0.1 M Tris-HCl pH 8.0; 0.2 M Magnesium chloride; and 20% (w/v) PEG 6000) from the PACT premier screen (Molecular Dimensions). In both cases, crystals were cryoprotected with the mother liquor supplemented with 20% (v/v) glycerol and frozen in liquid nitrogen before data collection at 100 K.

### Data processing, phase determination, refinement, and model building

Data were collected at the European Synchrotron Radiation Facility (ESRF) beamlines ID30A-3 and ID23-2 (DOI 10.15151/ESRF-DC-1823716253). The diffraction data were automatically processed by pipelines established at ESRF. The TEsup-1/dZfc3h1 diffraction dataset was processed by XIA2 (Winter *et al*, 2013) using DIALS (Beilsten-Edmands *et al*, 2020). The TEsup-1/Pih1D1 dataset was processed by the ESRF pipeline Grenoble Automatic Data Processing (GrenADES) using XDS,(Kabsch, 2010) pointless and aimless (Evans, 2011; Evans & Murshudov, 2013; Winn *et al*, 2011). For the TEsup-1/Pih1D1 structure, the phases were determined by molecular replacement using the corresponding domain of the AlphaFold model of TEsup-1 (AF-Q9VBX4-F1, residues 706-800) (https://alphafold.ebi.ac.uk/entry/Q9VBX4). The model was prepared with Phenix (process predicted model) to translate the pLDDT values to B factors and to remove flexible regions (Oeffner *et al*, 2022). Molecular replacement was performed with Phaser(McCoy *et al*, 2007) within Phenix (Liebschner *et al*, 2019). The model was automatically built with ModelCraft (Bond & Cowtan, 2022) within CCP4i2 (Potterton *et al*, 2018), manually completed with COOT (Emsley *et al*, 2010) and refined with phenix.refine(Afonine *et al*, 2012) and refmac5 (Kovalevskiy *et al*, 2018). Model quality was assessed using molprobity (Williams *et al*, 2018) and PDB-REDO (Joosten *et al*, 2012). For the TEsup-1/dZfc3h1 structure, the phases were determined using the TEsup-1 model from the refined TEsup-1/Pih1D1 structure with Phaser (McCoy *et al*., 2007) within Phenix (Liebschner *et al*., 2019). Model building, refinement, and validation were performed as described for TEsup-1/Pih1D1. Data collection and refinement statistics are shown (Data Table 1). Molecular graphics and analyses were performed with UCSF ChimeraX version 1.8 (Meng *et al*, 2023).

### ITC

ITC was performed using Microcal PEAQ-ITC calorimeter (Malvern Pananalytical). TEsup-1 PRD and the dZfc3h1 and Pih1D1 peptides were dialysed simultaneously in the following buffer: 20 mM Tris-HCl pH 7.5, 150 mM NaCl, 0.25 mM TCEP. For the TEsup-1 PRD/ dZfc3h1 combination, 39.6 μM TEsup-1 PRD and 396 μM dZfc3h1 peptide were used. Due to the low affinity of the Pih1D1 peptide for TEsup-1 RD, higher concentrations were required, with 382 μM TEsup-1 PRD and 3.8 mM Pih1D1 peptide. Every experiment included one injection of 0.4 μl, followed by eighteen injections of 2 μl dZfc3h1/Pih1D1 peptide into 200 μl of TEsup-1 PRD protein. All titrations were carried out at 25°C. The titrations of the dZfc3h1 and Pih1D1 peptides into the buffer served as negative controls. The data were processed with Microcal PEAQ-ITC analysis software v.1.41 and plotted and visualized using RStudio.

### Sequence analysis of CG31510/TEsup-1 and CG7065/TEsup-2

To support sequence similarity analyses of CG31510/TEsup-1 and CG7065/TEsup-2, their AlphaFold models were utilized to predict domain boundaries using Merizo in iterative mode. Four domains were identified in both proteins. In CG31510/TEsup-1, the predicted domains are located at 1-94 (confidence score: 0.953), 458-605 (0.997), 706-814 (0.612), and 848-949 (1.000). The first domain is unique to CG31510/TEsup-1, while the remaining three are shared with CG7065/TEsup-2. For CG706/TEsup-2, the predicted domains span positions 1-173 (0.883), 266-371 (0.762), 596-698 (1.000), and 834-903 (1.000), with the last domain being specific to CG7065/TEsup-2. Using the amino acid sequences of these domains as input for an HHpred search against the SMART v6.0 domain database, ZnF_U1/SM00451 domains were predicted in the first two structural domains of CG31510//TEsup-1 (positions 53-83, E=1.3e-06; 476-506, E=0.0066) and in the first structural domain of CG7065/TEsup-2 (positions 51-81, E=0.0089).

The second domain of CG31510/TEsup-1, which is shared with CG7065/TEsup-2, bears resemblance to U1-like zinc finger regions found in DZF proteins. Additionally, an SR-rich region was identified in CG7065/TEsup-2 between positions 436-547 using fLPS2 (2.688e-29). TEsup orthologs across the Athropoda phylum- crustaceans, springtails, insects, were obtained from OrthoDB v11. Members of the CG7065/CG31510 orthogroup 3616254at2759 were retained if they contained the three domains shared by TEsup1 and TEsup2- as determined by significant (E<0.01) sequence similarity in independent jackhmmer searches initiated with each domain sequence. An alignment of the PRD-domain was generated from this set of orthologs using mafft v.7.525, and a profile Hidden Markov Model was subsequently constructed using HMMER v3.4. A HMMER search with the PRD HMM profile against the Reference Proteomes hits CG7065-orthologs and also shows significant matches to FUBP-family proteins (e.g. A0A091NTP7/E=0.0013, A0A8K0ABI9/0.0071). FUBP3/Q96I24 is also the only significant hit in a profile search against the human proteome (E=0.0027).

### Ortholog identification and sequence analysis of CG4294/dZfc3h1

OrthoDB v11 was used to obtain a comprehensive set of one-to-one CG4294 orthologs from a wide range of insect taxa, including Diptera, Hymenoptera, Lepidoptera, Coleoptera, and Thripidae. After removing partial and duplicated sequences, 171 full-length proteins remained and were aligned using MAFFT v7.525. A profile Hidden Markov Model (HMM) of insect CG4294 proteins was then constructed using HMMER v3.4. ZC3H1_HUMAN was identified as the likely ortholog of insect CG4294, being the only significant hit with an E-value of 9e-31 in a HMMER search with the generated HMM profile against the human proteome (Reference Proteomes Release 2024_04). The first domain had a i-Evalue of 1.3e-07, covering residues 69-309 in the HMM and aligning with residues 18-400 of ZC3H1_HUMAN. The second domain had a i-Evalue of 1.1e-20, covering residues 471-1169 in the HMM and aligning with residues 399-1214 of ZC3H1_HUMAN. Reciprocal similarity was confirmed via an HHpred search using the ZC3H1_HUMAN region 835-1235 against the Drosophila proteome. Here, CG4294 was the only significant hit with an E-value of 5e-9, aligning with NP_611684.3 in the region 1043-1333 after three HHblits iterations with global realignment.

## Acknowledgements

We thank the VBCF core facilities (ProTech, NGS, VDRC) for support, and the IMBA Fly House for generating transgenic and CRISPR-edited fly lines. The GMI/IMBA/IMP Scientific Service units provided excellent assistance, especially the Mass spectrometry unit. We thank M. Madalinski for peptide synthesis and the Max Perutz Laboratories Monoclonal Antibody facility for creating the TEsup-1 and TEsup-2 hybridoma cell lines. We thank the beamline scientists from the European Synchrotron Radiation Facility (ESRF, Grenoble, France) for excellent support with data collection. We are grateful to V. Andreev, J. Batki, V. Loubiere, M. Matzinger, G. Dürnberger, B. Rafanel, A. Stark, M. Steiner and A. Tsarev for their help with preliminary experiments and discussions. Finally, we thank the entire Brennecke lab for their support.

## Funding

Work in the Brennecke laboratory was funded by the Austrian Academy of Sciences and the European Research Council (ERC-2015-CoG-682181). X-ray diffraction studies were carried out at ESRF, supported by the Austrian Science Fund (10.55776/I6110 and 10.55776/DOC177) to S.F. and a DOC fellowship from the Austrian Academy of Sciences to T.M.. C. Yu is supported by the VIP^2^ Post-Doc fellowship program as part of the EU Horizon 2020 research and innovation program (Marie Skłodowska-Curie grant No. 847548). U.H. was supported by a Marie Sklodowska-Curie fellowship (896416) and an EMBO long-term fellowship (ALTF_1175-2019). F.N. and J.P.M. are supported by Boehringer Ingelheim Fonds PhD Fellowships. E.B. is supported by a BBSRC DTP PhD Fellowship. F.K.T. is a Wellcome Trust Sir Henry Dale Fellow (206257/Z/17/Z), an EMBO YIP (5025), and is supported by the Human Frontier Science Program (CDA-00032/2018) and the Wellcome Trust (Discovery Award 317408/Z/24/Z).

## Author contributions

C.Y. and J.B. conceived and conceptualized the study. T.M. determined the X-ray structures of TEsup-1 PRD–peptide complexes and performed ITC assays under S.F.’s supervision. C.Y. carried out all molecular biology experiments, with contributions from U.H. and D.H. on AlphaFold predictions and peptide designs, J.P.M. on co-IPs, F.N. on PRO-seq experiments, and G.G. on recombinant protein purification. C.Y. and D.H. performed computational analyses, J.P.M. and M.N. carried out ortholog identification and phylogenetic analysis, J.S. helped with the initial transcriptional silencing assays with TEsup-1, L.T., J.B. and P.D. assisted with fly experiments. E.R. supervised the Mass Spectrometry analyses. F.K.T. and E.B. contributed to *P*-element experimental design, materials, and analyses. C.Y. and J.B. wrote the paper with input from F.K.T., T.M. and S.F.

## Declaration of interests

The authors declare that they have no competing interests.

## Data and materials availability

Diffraction data for the two crystal structures were collected at ESRF beamlines ID30A-3 and ID23-2 (DOI: 10.15151/ESRF-DC-1823716253). The coordinates and structure factors are available in the Protein Data Bank with accession codes 9G7L (TEsup-1/dZfc3h1) and 9G7U (TEsup-1/Pih1D1). All sequencing datasets (RNA-seq, ChIP-seq, PRO-seq, TT-seq, and small RNA-seq) have been deposited in the NCBI GEO archive (GSE274430, GSE274431, GSE274432, GSE274433, GSE313706, GSE313707, GSE313708, GSE313709, GSE313710) and can be accessed at this UCSC Genome Browser link (https://genome-euro.ucsc.edu/s/changwei/2025_TEsups). The mass spectrometry proteomics data have been deposited with the ProteomeXchange Consortium via the PRIDE repository (dataset identifier PXD054812). The *Drosophila melanogaster* genome dm6 version was used throughout this study. All fly strains established in this study are available from the VDRC stock center.

---

**Figure S1.**
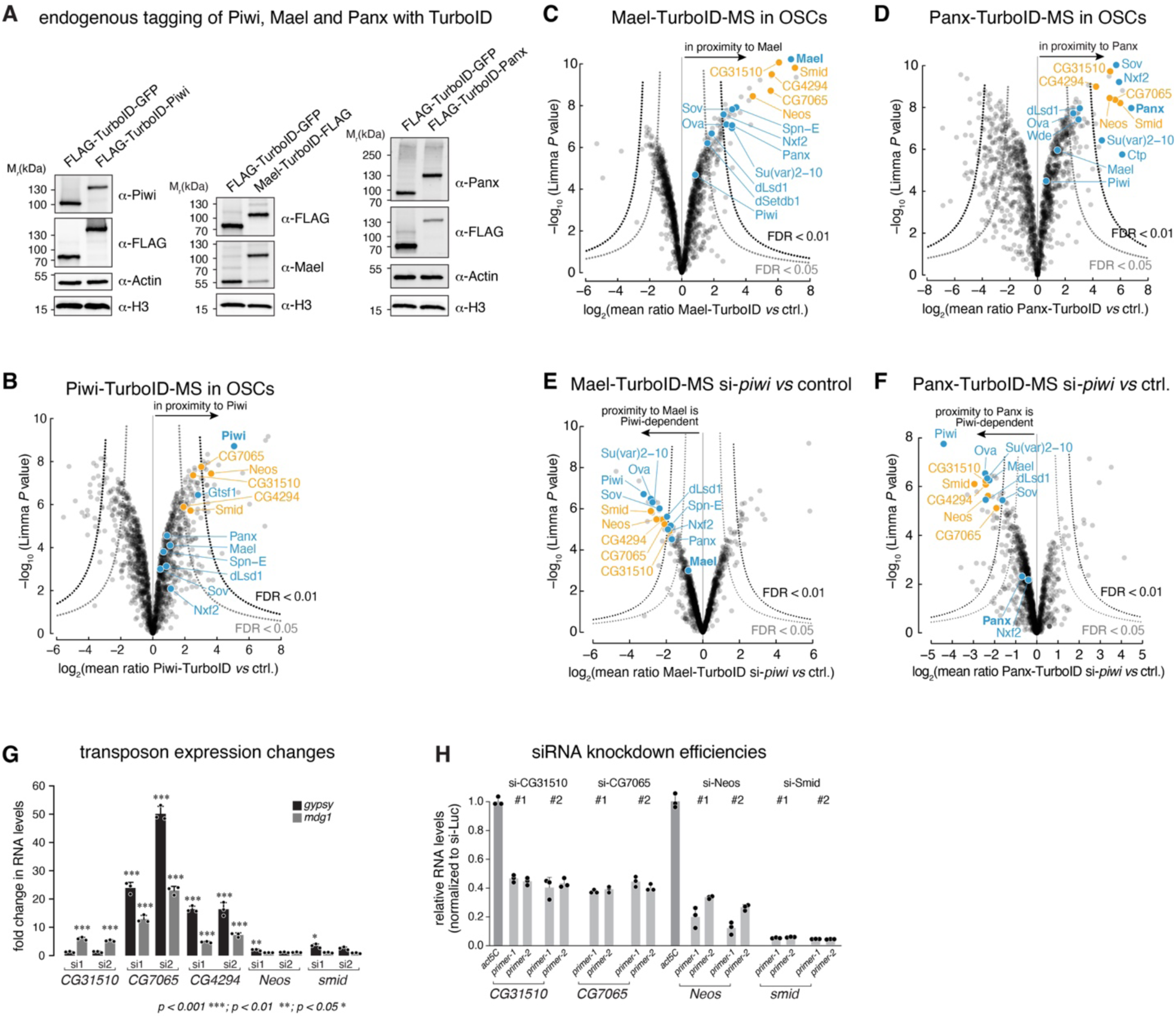
Piwi and cofactor proteomics in OSCs reveals novel silencing factors. **(A)** Western blot showing levels of TurboID-GFP-FLAG, FLAG-TurboID-Piwi, Mael-TurboID-FLAG and FLAG-TurboID-Panx in respective OSC lines. Blots were probed with antibodies against Piwi, Mael, Panx, FLAG, Actin, and Histone H3 (loading control). **(B)** Volcano plot showing protein enrichment from quantitative mass spectrometry comparing Piwi-TurboID with GFP-TurboID (n = 3 biological replicates; ctrl.: control). Known piRNA pathway factors are in blue, factors followed up in this study in orange. Significance was determined using limma (Smyth, 2004) and FDRs were calculated as in (Hein *et al*., 2015). **(C, D)** Similar to (B), for Mael-TurboID (C) and Panx-TurboID (D). **(E, F)** Volcano plot showing mass spectrometry results for Mael-TurboID (E) and Panx-TurboID (F) in Piwi-depleted versus control cells (n = 3 biological replicates). Significance, FDR, and labeling as in Fig. S1B. **(G)** RT-qPCR analysis of TE RNA levels after depletion of CG31510, CG7065, CG4294, Neos, or Smid (two independent siRNAs each), normalized to si-Luc control and *act5C* mRNA (mean of 3 replicates; error bars represent standard deviation; statistical significance was determined using Analysis of variance (ANOVA) test). **(H)** RT-qPCR analysis of *CG31510, CG7065, Neos* and *smid* knockdown efficiency following siRNA transfection (two different siRNAs each) using two qPCR amplicons (mean of 3 technical replicates; error bars represent standard deviation).

**Figure S2.**
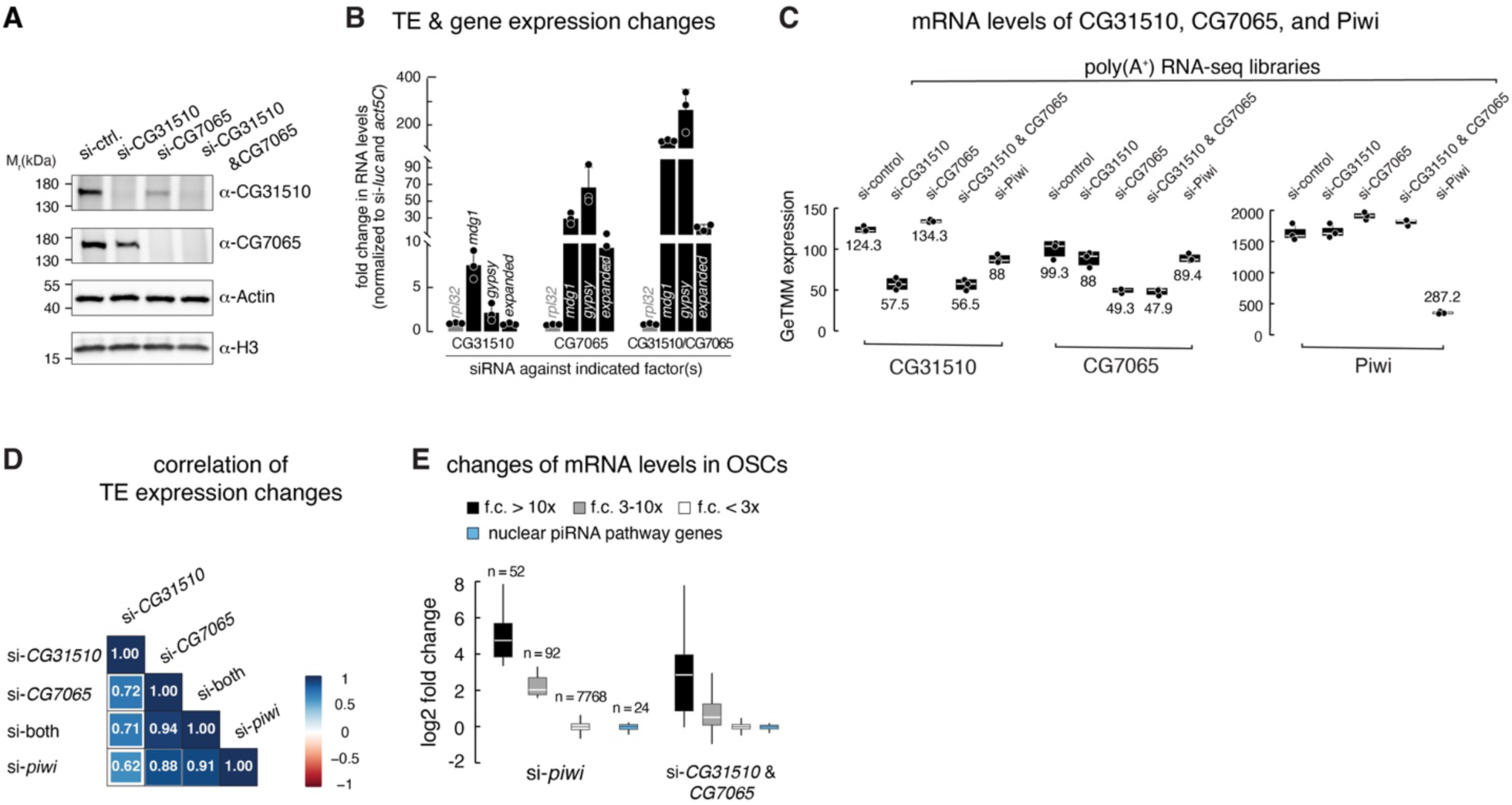
CG31510 and CG7065 are cofactors of Piwi-mediated silencing in OSCs. **(A)** Western blots showing siRNA-mediated depletion of CG31510 and/or CG7065 in OSCs, probed with antibodies against CG31510, CG7065, Actin, and Histone H3. **(B)** RT-qPCR analysis showing fold changes in steady-state RNA levels of indicated TEs and genes (*expanded* is a Piwi-repressed gene due to a close by *gypsy* insertion) in OSCs after depletion of CG31510 and/or CG7065, normalized to si-*Luc* control samples and *act5C* mRNA levels (mean of 3 biological replicates; error bars represent standard deviation). **(C)** Levels of *CG31510*, *CG7065* and *piwi* mRNA in indicated poly(A⁺) RNA-seq libraries. **(D)** Correlation of TE expression changes in OSCs depleted of CG31510, CG7065, both, or Piwi, based on poly(A⁺) RNA-seq analysis. **(E)** Boxplots showing mRNA level changes in Piwi-depleted and CG31510 & CG7065-depleted OSCs versus control. mRNAs are grouped into four categories: nuclear piRNA pathway-related genes (blue; n=24; Table S9), and three groups based on their fold changes in Piwi-depleted cells compared to control (fold change > 10 (black), 3-10 (grey), < 3 (white); GeTMM ≥ 5 in at least one dataset).

**Figure S3.**
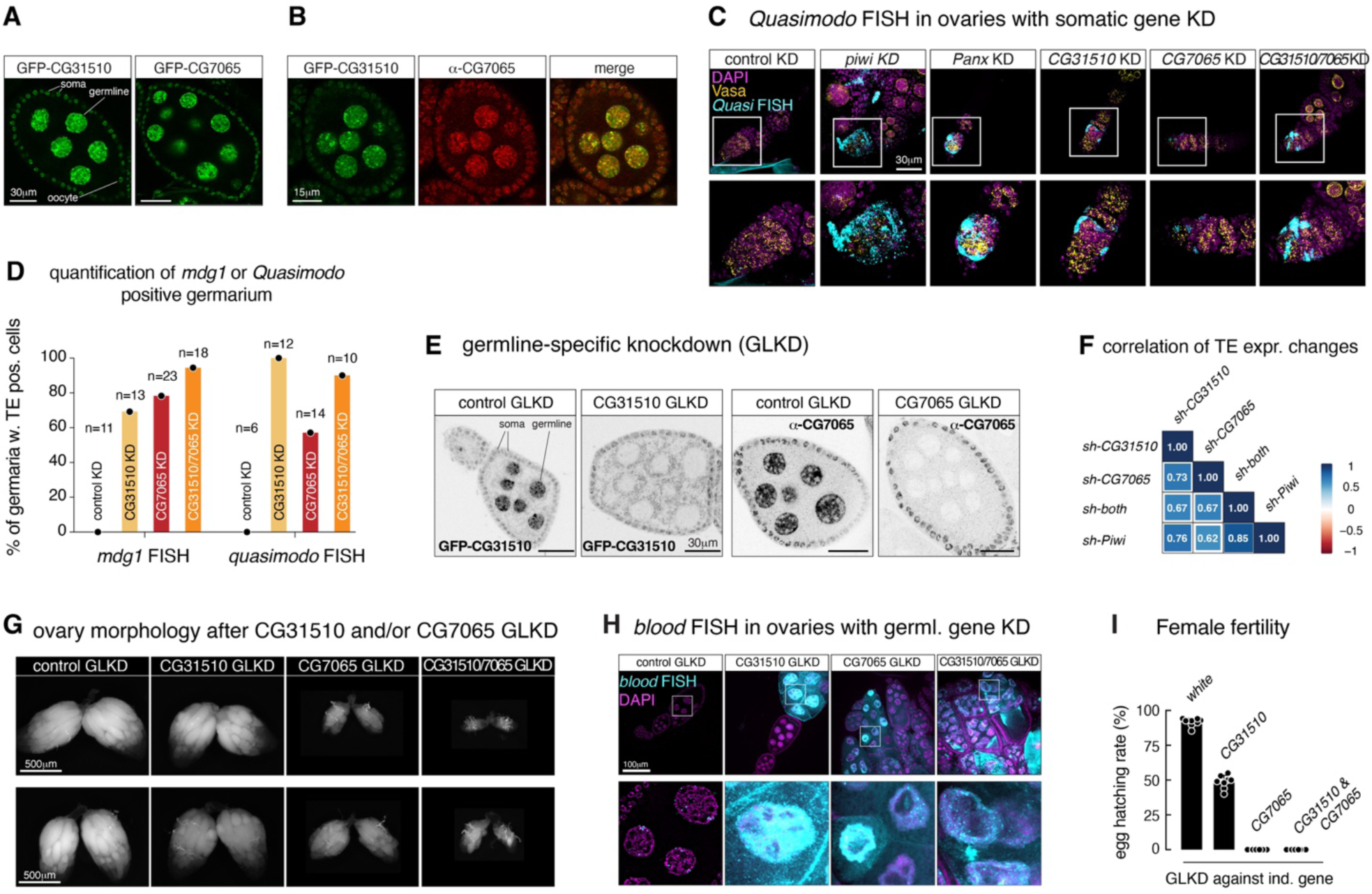
CG31510 and CG7065 are cofactors of Piwi-mediated silencing *in vivo*. **(A)** Confocal images of *Drosophila* egg chambers showing localization of GFP-tagged CG31510 and CG7065 in germline and somatic cells (scale bar: 30 µm). **(B)** Confocal images showing localization patterns of GFP-tagged CG31510 and endogenous CG7065 (assessed by anti-CG7065 immunofluorescence) in the same egg chamber (scale bar: 15 µm). **(C)** Confocal images of early oogenesis (germaria and stage 2 egg chambers) stained for DNA (DAPI), Vasa, and *Quasimodo* RNA (FISH). Enlarged views of boxed regions are shown below. **(D)** Quantification of germaria harboring *mdg1* or *Quasimodo* RNA FISH positive cells based on confocal images for Figure 1G and figure S3C (n = number of images analyzed). **(E)** Confocal images verifying shRNA-mediated depletion of CG31510 (GFP-tagged) and CG7065 (anti-CG7065 immunofluorescence) in the female germline. Germline-specific knockdown (GLKD) was achieved by MTD-Gal4-driven expression of shRNA hairpins (scale bar: 30 µm; remaining signal in the CG31510 knockdown sample corresponds to auto-fluorescence of mitochondria in the green channel). **(F)** Correlation of TE expression changes in ovaries depleted of CG31510, CG7065, both, or Piwi in the germline, based on poly(A⁺) RNA-seq analysis. **(G)** Bright-field images showing ovarian morphology from flies of indicated genotypes. Germline-specific knockdown (GLKD) was achieved by MTD-Gal4-driven expression of shRNA hairpins (scale bar: 500 µm). **(H)** Top: Confocal images showing *blood* RNA FISH signal in *Drosophila* egg chambers from flies of indicated genotypes (FISH signal: cyan; DAPI: magenta; scale bar, 30 µm). Bottom: Zoomed-in view of *blood* FISH signal in a germline nucleus from the top images (white boxes). **(I)** Hatching rates of eggs laid by females with indicated genotypes mated to wild-type males (mean of 8 biological replicates; error bars represent standard deviation).

**Figure S4.**
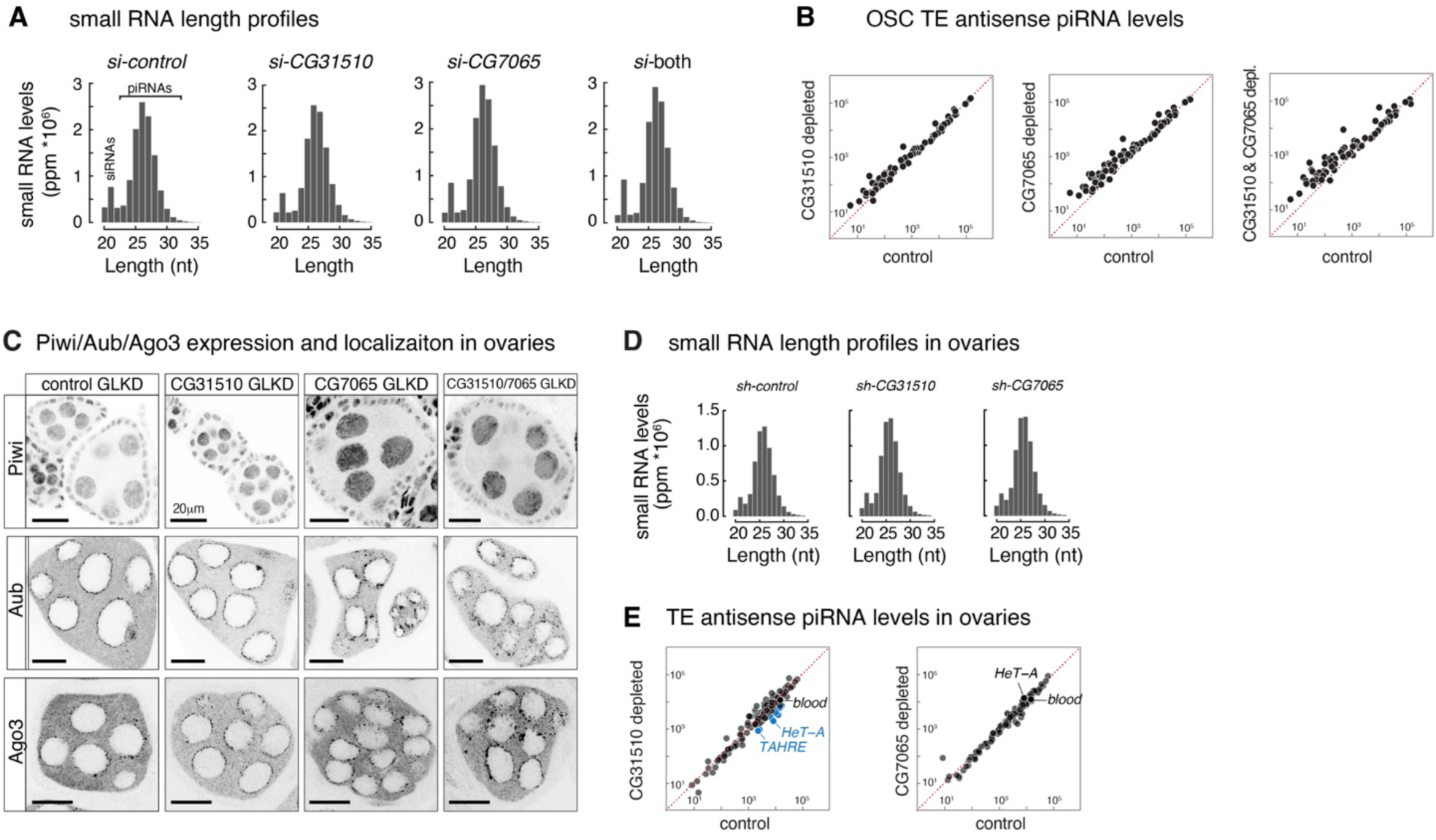
CG31510 and CG7065 do not affect overall piRNA biogenesis in OSCs and *in vivo*. **(A)** Bar graphs showing the levels and size distributions of small RNAs, normalized to one million miRNAs (ppm), in OSCs after siRNA-mediated depletion of indicated genes. **(B)** Scatter plots showing antisense piRNA levels targeting individual TE consensus sequences in the respective knockdown conditions compared to control. **(C)** Confocal images showing the levels and localization of the three PIWI proteins (Piwi, Aubergine, and Ago3) in *Drosophila* egg chambers with indicated genotypes (scale bar: 20 µm). **(D)** Bar graphs showing the levels and size distributions of small RNAs, normalized to one million miRNAs (ppm), in fly ovaries after shRNA-mediated depletion of indicated genes. **(E)** Scatter plots showing ovarian antisense piRNA levels targeting individual TE consensus sequences in the respective germline knockdown conditions compared to control.

**Figure S5.**
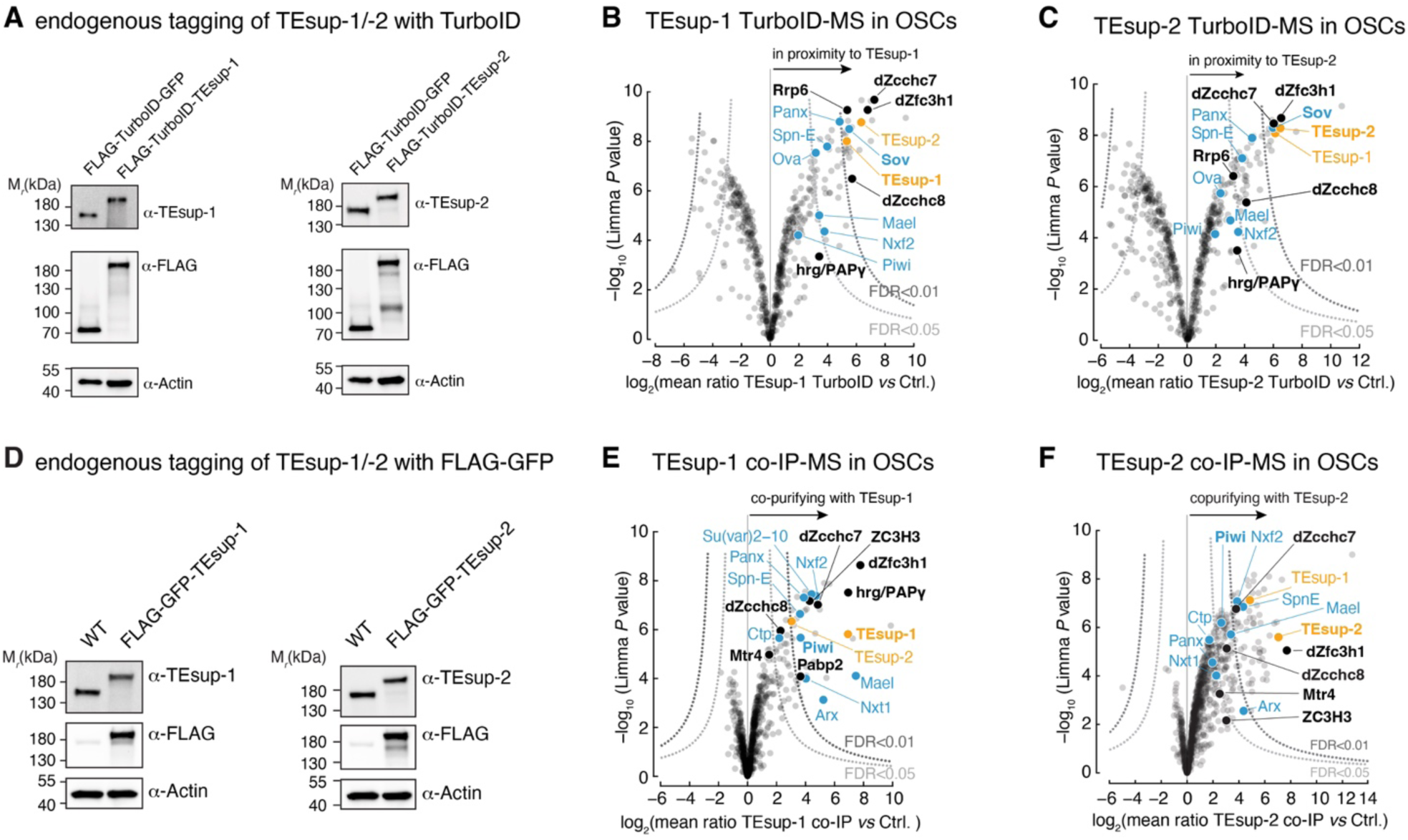
Protein interactors of TEsup-1 and TEsup-2. **(A)** Western blot showing levels of TurboID-GFP-FLAG, FLAG-TurboID-TEsup-1, and FLAG-TurboID-TEsup-2 in respective endogenously tagged cell lines. Blots were probed with antibodies against TEsup-1, TEsup-2, FLAG, and Actin (loading control). **(B)** Volcano plot displaying log2-fold enrichment of proteins identified by quantitative mass spectrometry, comparing TEsup-1-TurboID with GFP-TurboID control (n = 3 biological replicates; statistics as in Figure S1B). Known piRNA pathway factors are in blue, TEsup-1 and TEsup-2 in orange, and nuclear RNA exosome-related proteins in black. **(C)** Similar to (B), for TEsup-2-TurboID. **(D)** Western blots validating expression of endogenously tagged FLAG-GFP-TEsup-1 and TEsup-2 cell lines. Blots were probed with antibodies against TEsup-1, TEsup-2, FLAG, and Actin (loading control). **(E)** Volcano plot showing log2-fold enrichment and significance of proteins identified by quantitative mass spectrometry, comparing TEsup-1 co-immunoprecipitates with control samples (n = 3 biological replicates; color coding as in (B); statistics as in Fig. S4B). **(F)** Similar to (E), for TEsup-2 co-immunoprecipitation.

**Figure S6.**
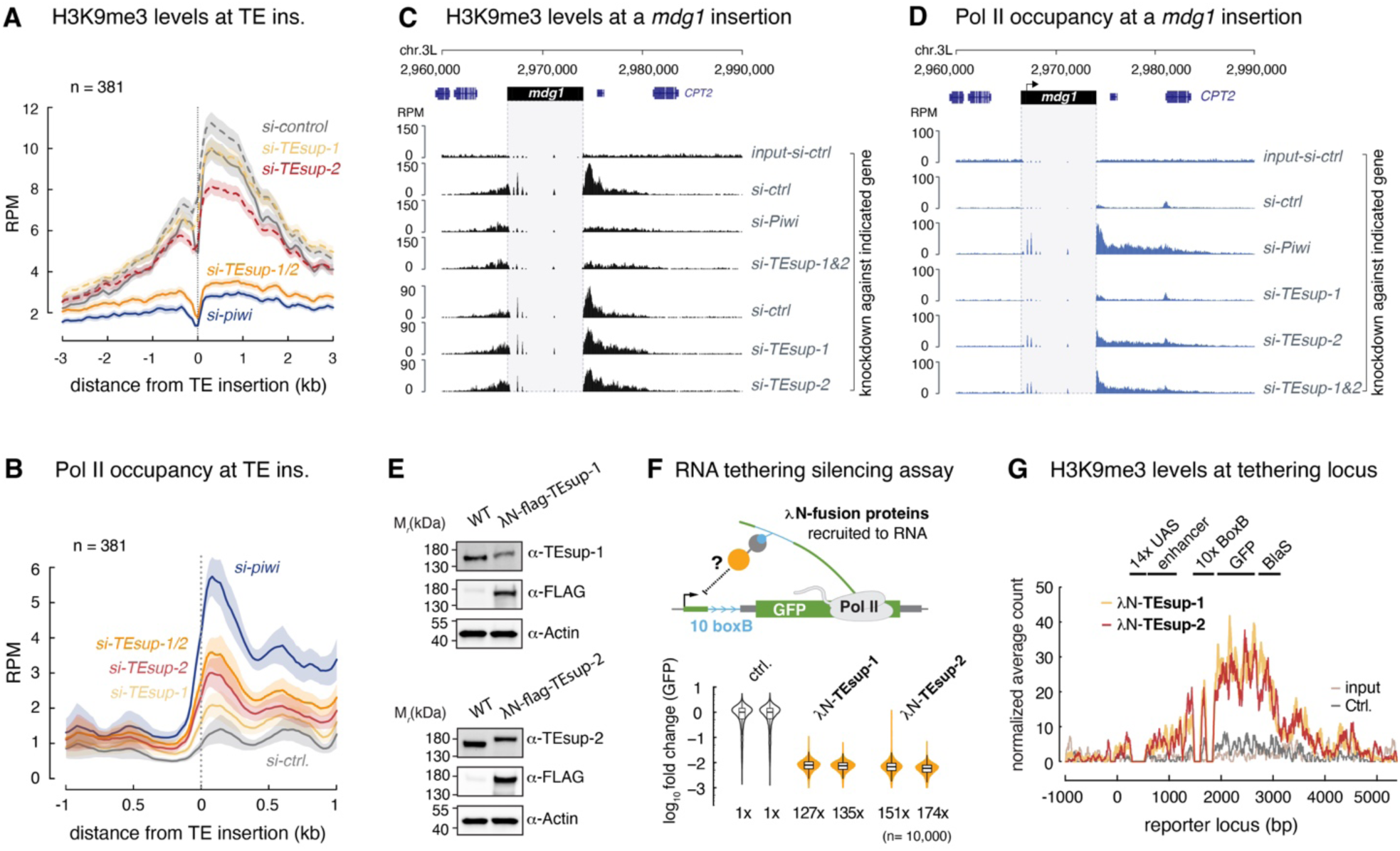
TEsup proteins promote Piwi-directed heterochromatin formation and transcriptional repression. **(A)** Metaplot of H3K9me3 levels around Piwi-regulated TE insertions (n = 381; vertical line) in control, TEsup-1/2, and Piwi-depleted OSCs (values represent means of n = 2 biological replicates; dashed line indicates experiment from a separated batch). **(B)** Metaplot of RNA Pol II occupancy around Piwi-regulated TE insertions (n = 381; vertical line) in control, TEsup-1, TEsup-2, TEsup-1/2, and Piwi-depleted OSCs (values represent means of n = 2 biological replicates). **(C)** UCSC genome browser tracks (30 kb window on chr. 3L) showing H3K9me3 occupancy around an *mdg1* insertion in OSCs after indicated siRNA knockdowns (average of n = 2 biological replicates; multi-mapping reads within the TE are not shown). **(D)** UCSC genome browser tracks showing RNA Pol II occupancy around an *mdg1* insertion after indicated siRNA knockdowns (average of n = 2 biological replicates; multi-mapping reads within the TE are not shown). **(E)** Western blots validating expression of endogenously tagged λN-FLAG-TEsup-1 and TEsup-2 in cell lines stably expressing a GFP reporter with an intron containing ten boxB hairpins. Blots were probed with antibodies against TEsup-1, TEsup-2, FLAG, and Actin (loading control). **(F)** Schematic of the λN-boxB GFP reporter silencing assay in OSCs (top), and violin plots of GFP reporter levels in cells stably expressing λN-tagged TEsup-1 or TEsup-2 (boxplots show median fold change; two biological replicates with 10,000 cells each; ctrl., GFP reporter cell line). **(G)** H3K9me3 levels at the reporter locus upon tethering of λN–TEsup-1, or λN–TEsup-2 and control (average profile of two biological replicates). Also shown are ChIP-seq input for λN–TEsup-1 (light grey) and reporter locus annotation.

**Figure S7.**
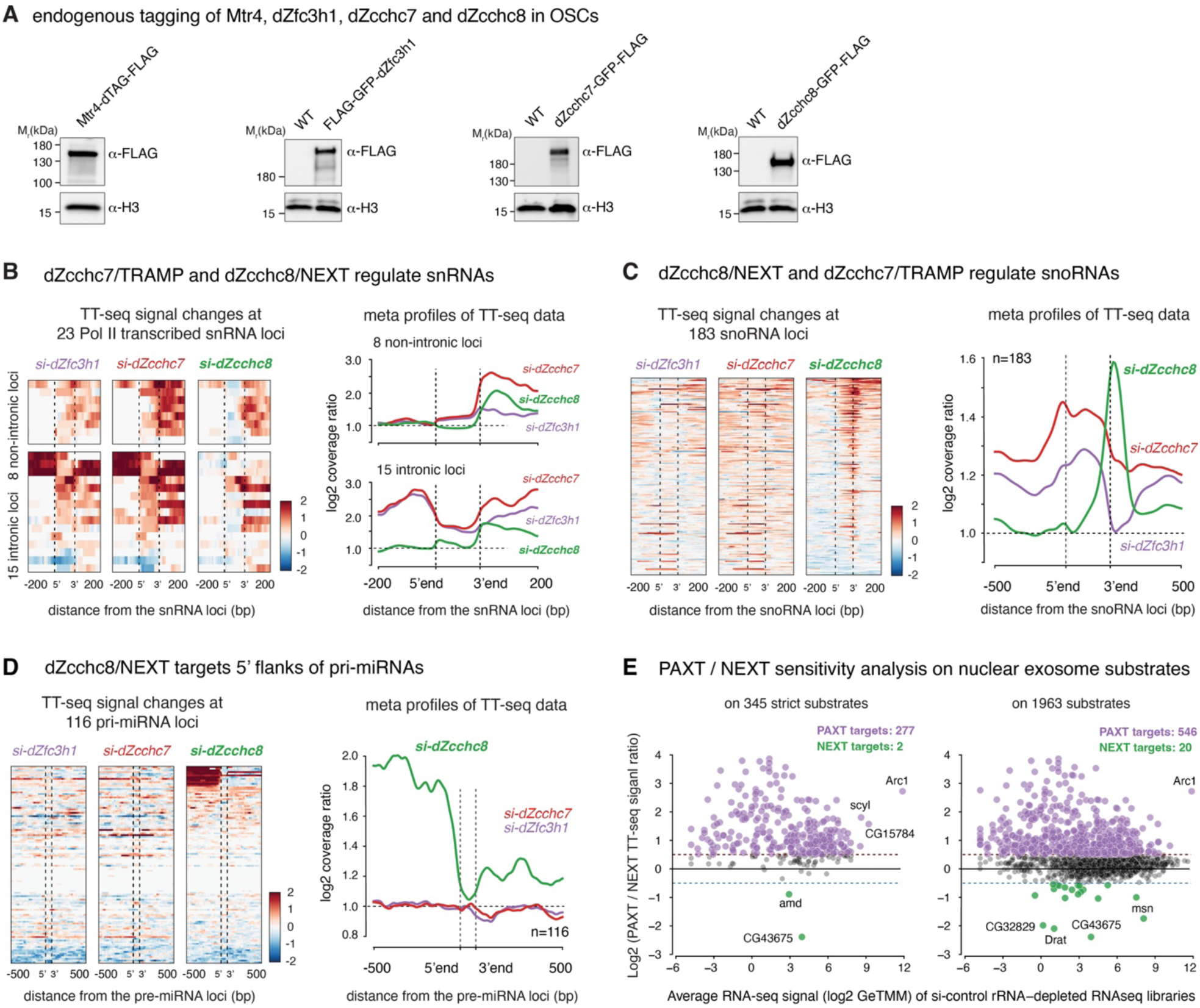
Characterization of nuclear RNA exosome adaptors in *Drosophila* OSCs. **(A)** Western blots verifying expression of endogenously tagged FLAG-dTAG-Mtr4 and FLAG-GFP-dZfc3h1, - dZcchc7, and -dZcchc8. Blots were probed with antibodies against FLAG and Histone H3 (loading control). **(B)** Left: Heatmap of log₂ transformed coverage of TT-seq signal at 23 Pol II transcribed snRNA loci (8 non-intronic loci, plus 15 intronic loci), after depletion of Zfc3h1 (purple), Zcchc7 (red), Zcchc8 (green) comparing to control. The snRNA area (5’ to 3’) was set to the mean length of the 23 Pol II transcribed snRNAs, extended 200bp to both upstream (5’) and downstream (3’) flanking regions. Right: Metaplots showing the mean values of log₂-transformed coverage of TT-seq signal, the ratio between experimental and control samples are displayed as for the heatmap (n = 3 biological replicates). **(C, D)** Similar to (B), for 116 snoRNA loci with detectable expression in OSCs (C), and for 116 pri-miRNA loci with detectable expression in OSCs (D) (n = 3 biological replicates). **(E)** PAXT and NEXT sensitivity analysis on nuclear exosome substrates from the TT-seq libraries, analysis was performed as described previously (Meola *et al*, 2016). Left: Analysis on 345 strict nuclear RNA exosome substrates (log₂ fold upregulation > 1 in all 6 nuclear RNA exosome related KD RNA-seq libraries). The y axis showing the log₂ PAXT:NEXT ratio (calculated as gene expression changes upon depletion of dZfc3h1 vs changes upon depletion of Zcchc8 from the TT-seq libraries). RNA targets with log₂ PAXT:NEXT ratios above 0.5 or below -0.5 were assigned as PAXT targets (purple) or NEXT targets (green), respectively. The x axis showing the log₂ average gene expression level (GeTMM) in control rRNA-depleted RNA-seq libraries. Each dot represents one nuclear RNA exosome substrate. Right: Similar to Left, for 1963 nuclear RNA exosome substrates (log₂ fold upregulation > 0 in all 6 nuclear RNA exosome related KD RNA-seq libraries).

**Figure S8.**
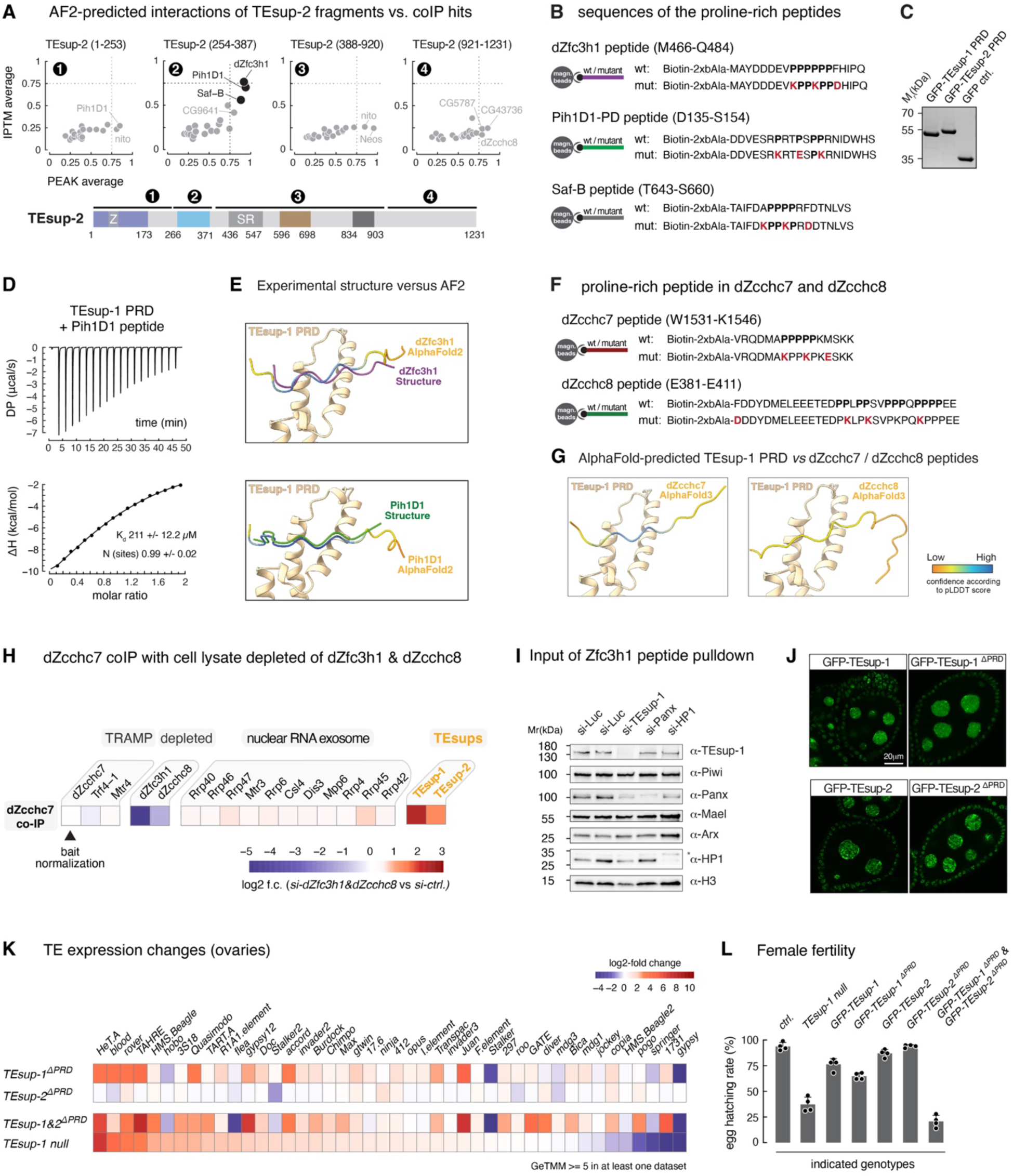
A proline-recognition domain in TEsup proteins binds exosome adaptors. **(A)** Results from an AlphaFold2 Multimer pairwise interaction screen between indicated TEsup-2 fragments (1–4) and significantly enriched TEsup-1&2 interactors (FDR < 0.05) identified by co-IPs. Direct interaction confidence is indicated by interface predicted Template Modeling (IPTM) and PEAK scores as in Fig. 3A. **(B)** Sequences of wildtype and mutant dZfc3h1, Pih1D1, and Saf-B proline-rich peptides with N-terminal biotinylation. **(C)** Coomassie-stained SDS–PAGE of recombinant GFP-tagged TEsup-1 PRD and TEsup-2 PRD. **(D)** ITC measurement of the interaction affinity between TEsup-1 PRD and the Pih1D1 peptide (n = 3 biological replicates). The interaction affinity (*K*_d_) and stoichiometry (N) represent the mean; the error corresponds to the standard deviation. **(E)** Top: Comparison of TEsup1-PRD–dZfc3h1 peptide structure with AlphaFold2 predicted model. Bottom: Comparison of TEsup1-PRD–Pih1D1 peptide structure with AlphaFold2 predicted model. **(F)** Sequences of wildtype and mutant dZcchc7 and dZcchc8 proline-rich peptides with N-terminal biotinylation. **(G)** Left: AlphaFold predicted model of TEsup-1-PRD–dZcchc7. Right: AlphaFold predicted model of TEsup-1-PRD–dZcchc8. **(H)** Heatmap showing log₂-fold enrichments of TRAMP subunits, nuclear RNA exosome subunits (black), and TEsup-1 & TEsup-2 (orange) in dZcchc7 co-IPs comparing wildtype OSC lysate to lysate from OSCs depleted of dZfc3h1 & dZcchc8 (Table S1). **(I)** Western blots showing the indicated proteins in the input samples for the peptide pulldown assays in Fig. 3I. **(J)** Confocal images of *Drosophila* egg chambers showing the localization and expression level of GFP-tagged TEsup-1, TEsup-1^ΔPRD^, TEsup-2, and TEsup-2^ΔPRD^ in germline and somatic cells (scale bar: 20 µm). **(K)** Heatmap of log₂ fold changes in TE RNA levels (poly(A⁺) RNA-seq) in ovaries from *TEsup-1^ΔPRD^*, *TEsup-2^ΔPRD^*, *TEsup-1&2^ΔPRD^*and *TEsup-1 null* mutant flies (n = 3 biological replicates). **(L)** Hatching rates of eggs laid by females with indicated genotypes mated to wild-type males (n = 4 biological replicates; error bars represent standard deviation).

**Figure S9.**
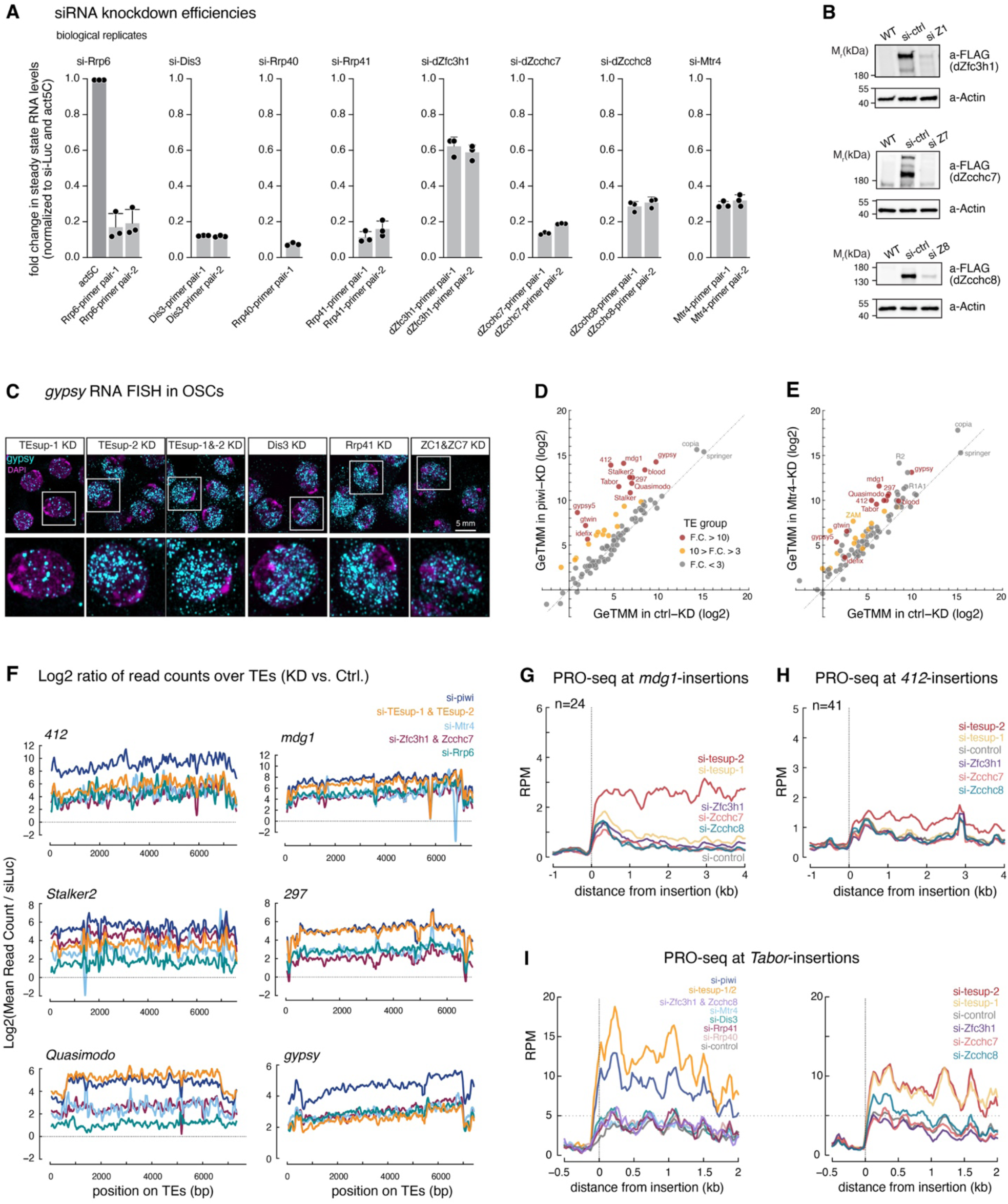
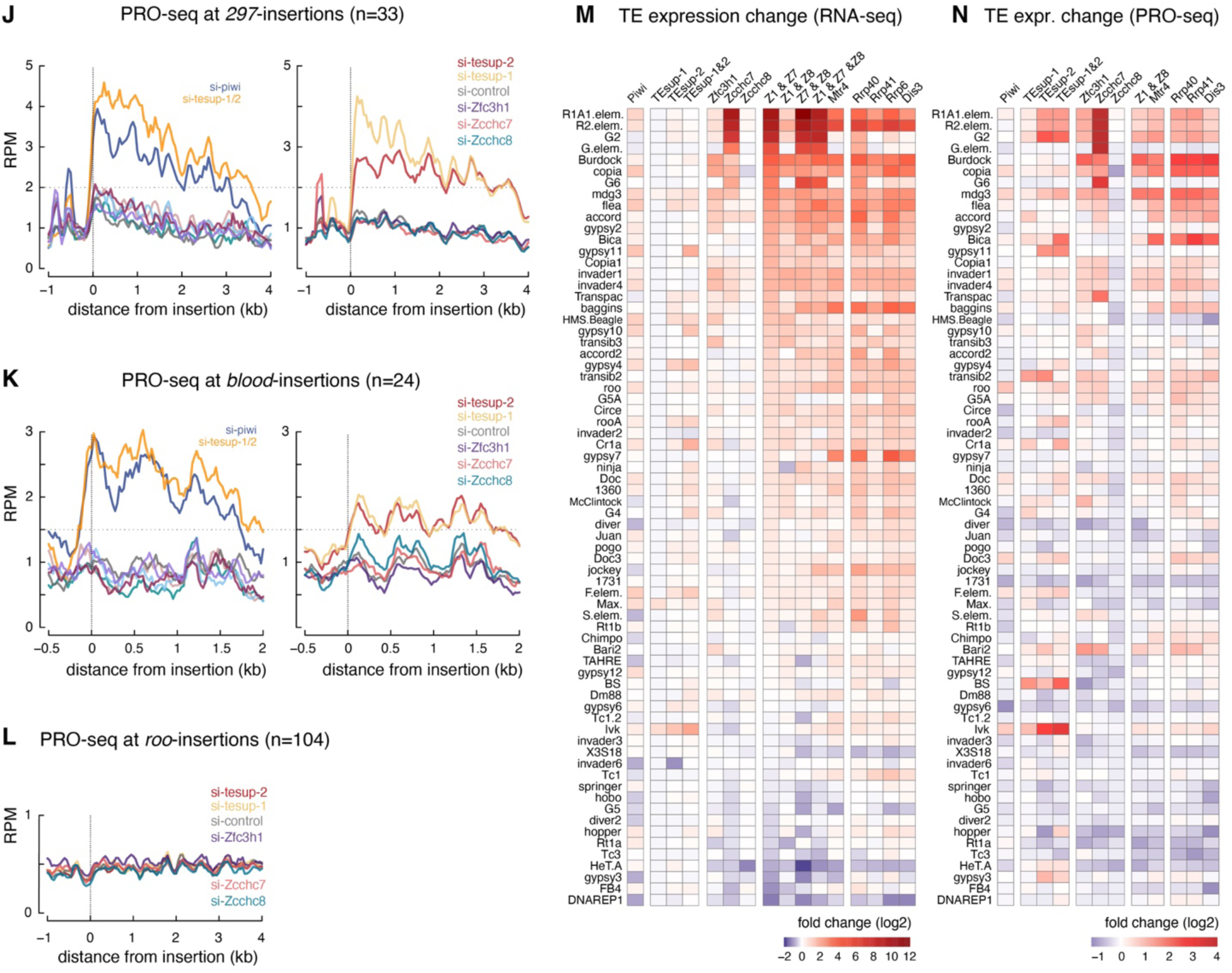
The nuclear RNA exosome is required for Piwi-mediated TE silencing. **(A)** RT-qPCR analysis probing siRNA knockdown efficiencies in OSCs, using one or two qPCR amplicons per gene. Data are presented as the mean of n = 3 biological replicates; error bars indicate standard deviation. **(B)** Western blot analysis showing FLAG-GFP-tagged dZfc3h1, dZcchc7, and dZcchc8 levels after indicated knockdowns. Blots were probed with antibodies against FLAG and Actin (loading control). **(C)** Confocal images of OSCs depleted for indicated factors by siRNAs and stained for DNA (DAPI) and *gypsy* RNA (FISH). Enlarged views of boxed regions are shown below. Same set as in Fig. 4B. **(D)** Scatter plot showing steady-state RNA levels of annotated TEs in Piwi-depleted OSCs versus control (ribo-zero RNA-seq, n = 3 biological replicates). TEs are grouped based on fold change (group I: >10; group II: 3-10; group III: <3). **(E)** Scatter plot showing steady-state RNA levels of annotated TEs in Mtr4-depleted OSCs versus control (ribo-zero RNA-seq, n = 3 biological replicates), similar to (D). **(F)** Metaplots showing the mean values of log₂-transformed coverage of RNA-seq signal for indicated TEs; the ratio between depletion condition and corresponding control is displayed (n = 3 biological replicates). **(G-L)** Metaplots of PRO-seq signal at genomic regions flanking indicated TE insertions following depletion of indicated genes (*mdg1, 412, tabor*, *297*, and *blood* are piRNA repressed TEs, while *roo* is not piRNA repressed). **(M)** Heatmap showing log₂-fold change of steady-state RNA levels for group III TEs (not piRNA repressed) in OSCs with indicated siRNA knockdowns (ribo-zero RNA-seq, n = 3 biological replicates). **(N)** Heatmap showing log₂-fold change of nascent transcriptional output for group III TEs (not piRNA repressed) in OSCs with indicated siRNA knockdowns (PRO-seq, n = 2 biological replicates).

**Figure S10.**
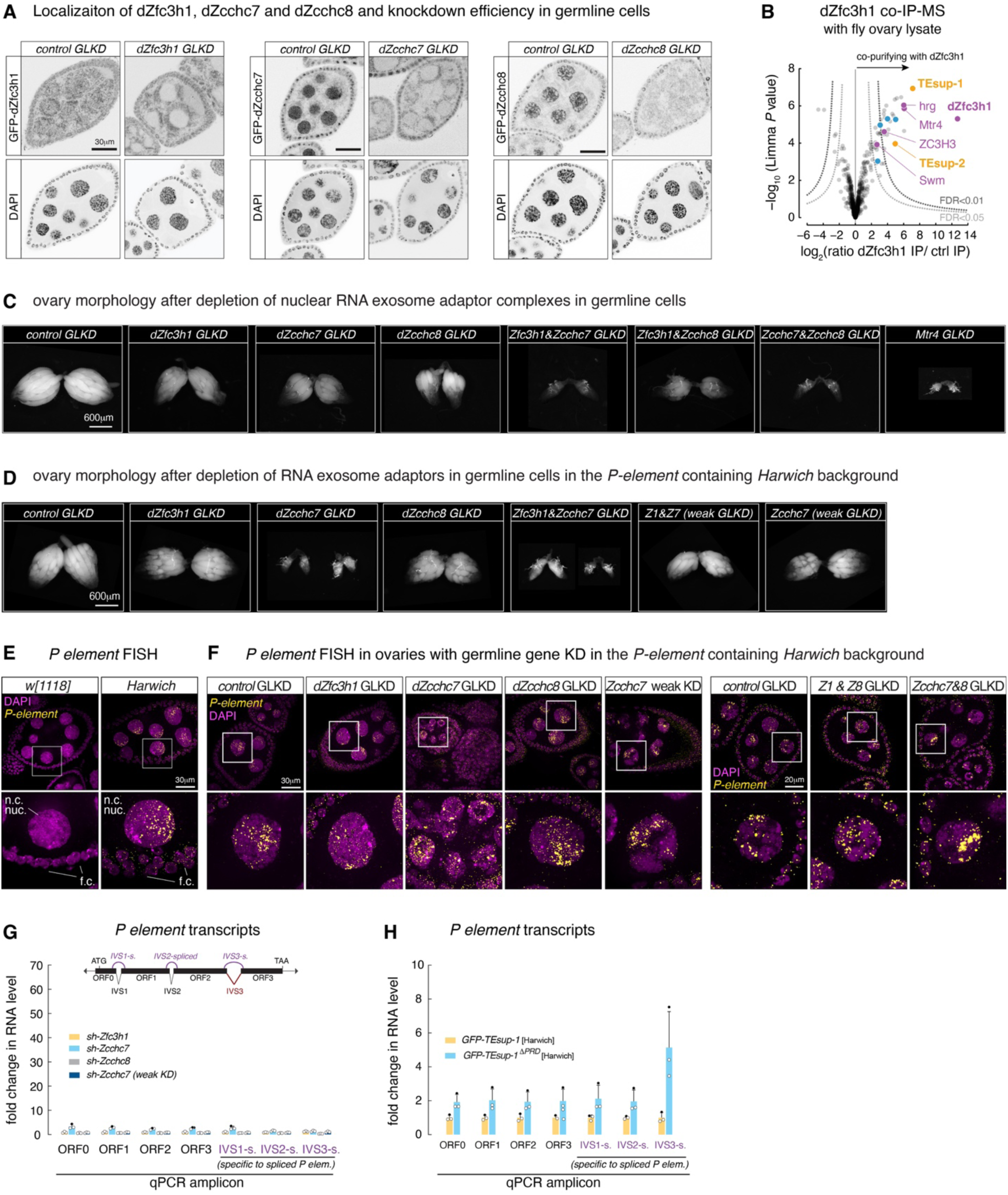
TEsup proteins and the nuclear exosome are critical for Piwi-mediated *P-element* repression in the *Drosophila* germline. **(A)** Confocal images verifying shRNA-mediated depletion of dZfc3h1 (GFP-tagged), dZcchc7 (GFP-tagged) and dZcchc8 (GFP-tagged) in the female germline. Germline-specific knockdown (GLKD) was achieved by MTD-Gal4-driven expression of shRNA hairpins (scale bar: 30 µm; cytoplasmic signal represents autofluorescence from mitochondria in the green channel). **(B)** Volcano plot showing log₂-fold enrichment and significance of proteins identified by quantitative mass spectrometry, comparing dZfc3h1 co-immunoprecipitates with control samples from *Drosophila* ovary lysates (n = 3 biological replicates; Table S1). Highlighted are subunits of PAXT (purple), TEsup proteins (orange), and Piwi silencing factors (blue). **(C)** Bright-field images showing ovarian morphology from flies of indicated genotypes. Germline-specific knockdown (GLKD) was achieved by MTD-Gal4-driven expression of shRNA hairpins (scale bar: 600 µm). **(D)** Similar to (C), for the indicated germline-specific knockdowns in the *P-element* containing *Harwich* background. **(E)** Confocal images of egg chambers from the *w[1118]* laboratory strain (lacking *P*-elements) or the *Harwich* strain (carrying multiple *P*-elements) stained for DNA (DAPI) and *P*-element RNA (FISH). Note nuclear export of *P*-element transcripts in somatic follicle cells (f.c.) but not germline nurse cells (n.c.). Enlarged views of boxed regions are shown below. **(F)** Confocal images of egg chambers from *Harwich* flies depleted for indicated factors in the germline using *nanos*-GAL4 driven expression of shRNA hairpins, stained for DNA (DAPI) and *P*-element RNA (FISH). Enlarged views of boxed regions are shown below. **(G)** RT-qPCR analysis showing fold changes in *P*-element RNA levels, using primers detecting the four coding exons (ORF0-3) and primers specific for spliced products of the three introns (IVS1s.-3s.), in ovaries depleted of indicated factors in the germline using *nanos*-GAL4 driven shRNA transgenes. Data are normalized first to *rpl32* mRNA level and then to sh-*white* control (mean ± s.d., n = 3 biological replicates). **(H)** As in (G), for ovaries from *GFP-TEsup-1^ΔPRD^* flies and the corresponding *GFP-TEsup-1* wildtype control, both carrying *P*-elements from the *Harwich* background and reared at 18 °C.

**Data Table 1.** Data collection and refinement statistics for TEsup-1 PRD–peptide crystal structures. *Values in parentheses are for the highest-resolution shell #RMSD: Root Mean Square Deviation

RMSD: Root Mean Square Deviation
|  | TEsup-1/dZfc3h1 | TEsup-1/Pih1d1 |
| --- | --- | --- |
| **PDB code** | 9G7L | 9G7U |
| Space group | C 1 2 1 | I 2 2 2 |
| Cell dimensions |  |  |
| *a*, *b*, *c* (Å) | 121.57, 48.63, 66.58 | 41.39, 75.09, 112.01 |
| $\alpha$, $\beta$, $\gamma$ (°) | 90.0, 113.04, 90.0 | 90.0, 90.0, 90.0 |
| **Data Collection** |  |  |
| Resolution range (Å) | 61.27–1.90 | 62.37–2.10 |
| No. of unique reflection | 28147 | 10539 |
| $R_{\text{merge}}$* | 6.7 (135.0) | 9.4 (167.9) |
| $R_{\text{pim}}$* | 4.1 (81.7) | 2.7 (46.9) |
| $I / \sigma I$* | 8.0 (0.7) | 13.7 (1.5) |
| Completeness (%)* | 99.0 (99.2) | 99.9 (99.5) |
| Multiplicity* | 3.6 (3.7) | 13.3 (13.7) |
| $CC_{1/2}$* | 99.7 (54.4) | 99.9 (88.4) |
| **Refinement** |  |  |
| Resolution range | 61.27–1.90 (1.97–1.90) | 37.54–2.10 (2.31–2.10) |
| Reflections used in refinement | 28023 | 10521 |
| $R_{\text{work}} / R_{\text{free}}$ (%) | 21.5 / 23.3 | 25.5 / 25.9 |
| Wilson B-factor (Å<sup>2</sup>) | 38.4 | 47.6 |
| Average B-factors (Å<sup>2</sup>) | 51.0 | 64.4 |
| No. of atoms | 2336 | 1082 |
| Proteins | 2239 | 1066 |
| Ligands | 24 | 6 |
| Solvent | 73 | 10 |
| **Stereochemistry** |  |  |
| RMSD<sup>#</sup> bond lengths (Å) | 0.002 | 0.004 |
| RMSD<sup>#</sup> bond angles (°) | 0.59 | 0.70 |
| Ramachandran favored (%) | 99.2 | 96.7 |
| Ramachandran outliers (%) | 0 | 0 |

## Notes

### Competing Interest Statement

The authors have declared no competing interest.

